# Closing the phenotyping gap: non-invasive electromagnetic induction enables high-throughput screening of crop root function in the field

**DOI:** 10.64898/2026.10.28.762415

**Authors:** Cheng Qian, Daniel Rodriguez, Fernando A. Monteiro Santos, Alison Kelly, Peter de Voil, David Jordan, Dongxue Zhao

## Abstract

Crop root systems play a critical role in water uptake and yield improvement under drought. However, a lack of methods to measure root system function at scale, particularly in the field, poses significant challenges to explore them in crop improvement programs. To address this bottleneck, we introduce a novel approach to directly screen water uptake and root activity in the field through integrating electromagnetic induction (EMI) sensing, inversion modelling and machine learning. This study was conducted as an on-farm field experiment comprising a factorial combination of 13 sorghum genotypes and three row spacings with three replications. Time-lapse EMI surveys were conducted from flowering to maturity using a customised DualEM-21S sensing system. Artificial neural network models were developed to predict 3D soil moisture (θ) using depth-specific true soil electrical conductivity estimated from EMI inversion algorithms, and the prediction accuracy was evaluated. Crop water use and functional root traits were derived from changes in θ profiles between successive EMI surveys using cubic splines fitted within a linear mixed model framework. Results indicate that dynamics of θ can be predicted with high accuracy (R^2^ = 0.83). There was a significant positive correlation between the EMI-derived crop water use and root length density at the depths without water stress. The functional root traits were significantly affected by growth stages, genotypes, row spacings, and their interactions. In general, narrower row spacing resulted in deeper rooting depth by flowering, which was associated with a larger canopy and higher grain number. In contrast, wider row spacing promoted greater post-flowering root growth, leading to increased post-flowering water use and, consequently, larger grain weight. Overall, our approach enables rapid characterisation of variation in crop water use, rooting depth and root growth in the field. It provides a valuable framework for understanding the dynamic interactions between root function and yield traits across variable field environments, with potential to accelerate genotype selection and optimise crop design in breeding and agronomy programs.

## 1. Introduction

Sorghum is the fifth-largest grain crop grown globally in areas with marginal rainfall and is a staple food for about 500 million people in Africa and Asia (FAO, 2023). With rapid global population growth and instability in seasonal rainfall patterns, there is a greater demand to improve plant traits that maximise sorghum yield in drought-prone environments. Improved yields in such environments are closely related to enhanced post-flowering water use and thus to increased water availability at flowering and/or increased water accessibility during grain filling (Van Oosterom et al., 2011). Under water-limited conditions, the temporal patterns and total amount of water use by the crop are largely determined by the root system’s capacity to grow to deeper soil depths and efficiently uptake available water at those depths. Genotypic differences in these root traits and their functional implications for water uptake and yield improvement under drought have long been recognised in cereal crops (Mace et al., 2012), while opportunities to explore them in crop improvement programs have been largely overlooked. This is mainly due to the difficulty in measuring root traits at scale, particularly in field environments.

So far, methods for screening roots have focused on 2D or 3D visualisations of root architectural (e.g., nodal root angle) or morphological (e.g., root hairs) traits, usually on young plants grown in aeroponics or small containers filled with artificial substrates (Richard et al., 2015). Although these methods are important, weak correlations have often been reported between seedling root traits and the field performance of mature plants. In the field, root phenotyping has mainly relied on excavation or soil coring techniques (Wasson et al., 2014), which are invasive and difficult to scale to the large numbers of genotypes required in breeding programs. Recently, geophysical methods such as ground penetrating radar (GPR), electrical resistivity tomography (ERT), and electromagnetic induction (EMI), have been proposed for non-invasive screening of root activity in the field by inferring water extraction patterns from soil water depletion profiles (Zhao et al., 2022). GPR can be used to detect coarse roots (diameter > 2mm) and is subject to strong signal attenuation in highly conductive soils. ERT measures soil electrical resistivity, which is sensitive to soil moisture, reflecting the entire root-zone soil under plants, reducing the influence of small-scale soil heterogeneity. However, ERT requires direct electrode contact with the soil, limiting its throughput compared with EMI sensors.

EMI sensors measure soil apparent electrical conductivity (EC_a_, mS/m) in a contactless way, enabling rapid collection of dense spatial datasets across large areas. Since soil properties affecting EC_a_ such as clay content, are relatively stable over time, changes in time-lapse EC_a_ measurements in non-saline crop fields can largely be attributed to changes in soil water content. EC_a_ represents a depth-weighted average of soil electrical conductivity over the sensor’s depth of exploration. Geophysical inversion modelling can be used to estimate depth-specific true soil electrical conductivity (σ) from EC_a_, which can then be converted to soil volumetric water content (θ) using an empirical model calibrated against ground-truth θ measurements. Quasi-2D inversion (Monteiro Santos 2004), which applies lateral smoothness constraints to produce spatially coherent estimates of σ, has been successfully used to map θ dynamics. Recently, several studies have demonstrated that a newly developed spatiotemporal inversion can improve estimates of θ and salinity dynamics by accounting for both temporal continuity and spatial variation in time-lapse EC_a_ data (Huang et al., 2016), particularly when θ varies continuously over time. However, this approach has not yet been evaluated for quantifying plot-scale θ dynamics under complex on-farm conditions, such as episodic rainfall and active crop transpiration. Furthermore, few calibration algorithms have been applied to small-scale field experiments to test hypotheses in agricultural research.

Field trials often show large spatial variation, which can confound the true genetic signal in crop phenotyping studies and add uncertainty to treatment comparisons in agronomic experiments (Gilmour et al., 1997). This variation arises from both the underlying field conditions (e.g., soil texture, sub-soil constraints, moisture, nutrition), and the management practices applied to the field trials, and it is necessary to model this spatial variation for accurate estimation of treatment and/or genotypic effects. Statistical models can be used to partition the effect of genotype (G), environment (E) and applied management (M), together with the two-way and three-way interactions of G×E×M combinations, and simultaneously account for experimental error and spatial dependence (Gilmour et al., 1997). These advanced statistical models are crucial for analysing root traits and crop water use, given the high spatial variability of soil. Moreover, understanding the relationships among above- and below-ground crop traits is important for identifying opportunities for genetic improvement in water-use efficiency. The potential value of the root traits should be examined through their relationships with water use, yield, and yield components, allowing for indirect selection and for assessing the potential benefits and trade-offs of selecting for root traits on agronomically important traits, e.g., yield.

Here, we applied an EMI-based root phenotyping approach to an on-farm experiment with 13 sorghum hybrids grown at three row spacings (0.5, 1 and 1.5 m). The objectives were to (i) determine which EMI inversion algorithm (quasi-2D or spatiotemporal inversion) is more robust for estimating soil water dynamics in the field; (ii) untangle G×M effects on sorghum root water use and root growth; and (iii) evaluate the functional relationships among root traits, crop water use, grain yield and yield components.

## 2. Materials and methods

### 2.1 Site description and experiment design

An on-farm trial was conducted at Tummaville (27°50’56.3” S 151°27’33.6” E), Qld, Australia (Fig. 1), during the 2023-2024 summer season (December to March). Treatments included 13 sorghum hybrids (coded A-M) sown at three row spacings (0.5, 1.0 and 1.5 m). The three row spacings were used to generate contrasting levels of intraspecific competition (Pielou, 1962). A split-plot design with three replications was used, in which row spacings were assigned to main plots, and hybrids were allocated to subplots. In total, there were 117 subplots, each 7m long by 4m wide.

**Fig. 1.**
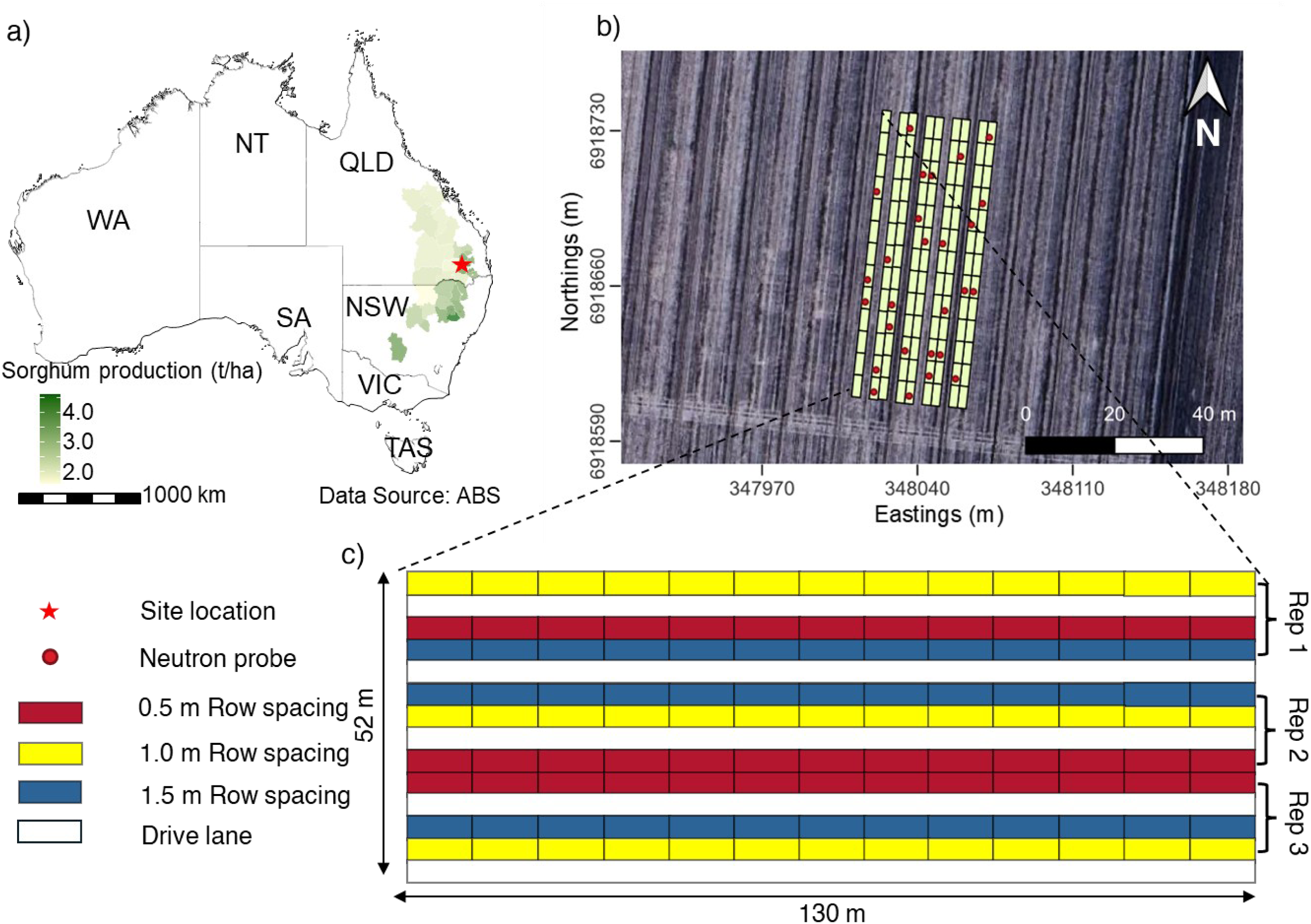
Overview of the experiment site and design. (a) Geographical distribution of sorghum production in Australia (ABS 2022–2023) and location of the experimental site; (b) experiment layout, with polygons representing subplots and red dots indicating locations of neutron moisture meter (NMM) access tubes; and (c) layout of the three row-spacing treatments and replications.

Soil at the site is classified as a uniform, self-mulching black Vertosol with clay content of above 60% (Fig. S1). An automatic weather station was installed on-site to record air temperature and rainfall throughout the growing season (Fig. 2 and S2). Total rainfall between sowing and flowering was 226 mm, equivalent to 128% of the long-term average (1950–2024) for the corresponding period, indicating wetter-than-average conditions during this period.

**Fig. 2.**
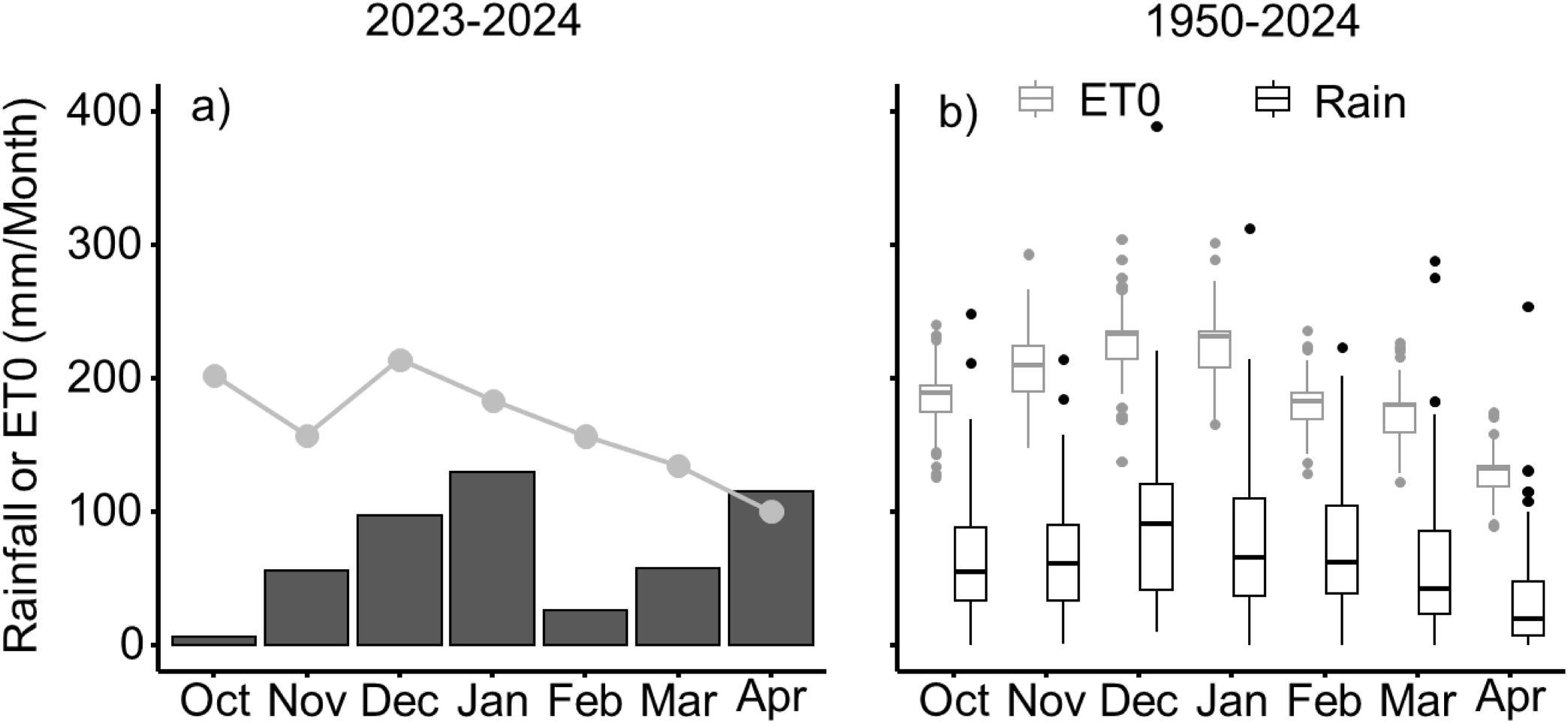
Seasonal rainfall and evaporative demand at the experiment site. (a) Monthly rainfall (bars) and reference evaporation (ET0, solid line) during the 2023–2024 summer growing season; (b) long-term rainfall (black) and ET0 (grey) distributions for 1950–2024. Boxes indicate the 25th–75th percentiles, central lines indicate medians, and whiskers extend to the 10th and 90th percentiles. Values outside this range are shown as points. Long-term climate data were obtained from https://www.longpaddock.qld.gov.au/silo.

The trial was sown on 12 December 2023 at a constant plant population density of 6 plants/m^2^. Liquid nitrogen fertiliser (Easy N, Incitec Pivot Fertilisers Ltd.) was applied at sowing to prevent nitrogen deficiency. Weeds, pests and diseases were managed as needed.

### 2.2 Time-lapse EMI surveys

A DUALEM-21S sensor (DualEM, Milton, Ontario, Canada) was mounted in a PVC pipe and towed by a vehicle to measure soil EC_a_ (mS/m) across the site, following Zhao et al. (2022). The sensor simultaneously measures EC_a_ over four nominal depths of exploration: 0–0.5, 0–1, 0–1.5, and 0–3 m. EMI surveys were conducted at 1–2-week intervals from flowering to maturity (Fig. S2).

To validate the EMI predictions, neutron moisture meter (NMM) measurements were collected in 27 plots across the field (Fig. 1b) on the same days as the EMI surveys. Measurements were taken at seven depth intervals spanning 0–2.1 m, including 0–0.3, 0.3–0.6, 0.6–0.9, 0.9–1.2, 1.2–1.5, 1.5–1.8 and 1.8–2.1 m. NMM measurements were collected on five occasions: three during flowering (14, 21 and 27 February) and two during grain filling (6 and 19 March).

### 2.3 EMI inversions

The EC_a_ data were inverted in EM4Soil software (V-4.8) (EMTOMO, 2025) using quasi-2D inversion (Monteiro Santos 2004) and spatiotemporal inversion (Huang et al. 2016) algorithms, respectively. In brief, the quasi-2D inversion estimates depth-specific true soil electrical conductivity (σ) using spatial constraints between neighbouring locations. Spatiotemporal inversion extends this approach by incorporating the modification proposed by Kim et al. (2011), applying regularisation in both space and time.

An iterative process starting from a uniform model was applied in EM4Soil to obtain a final model, with its response fitting the observed data in a least-squares sense. A forward modelling subroutine based on the full solution (FS) of the Maxwell equations (Wait, 1982) was used in both inversion algorithms given the large EC_a_ values at the site (> 100 mS/m). EM4Soil provides two regularization models, S1 and S2 (Sasaki, 2001, 1989). In the S1 option, model parameters are updated at each iteration using the following equation

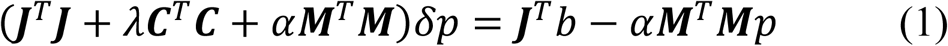

In the S2 option, the model parameters at each iteration are adjusted based on the following equation

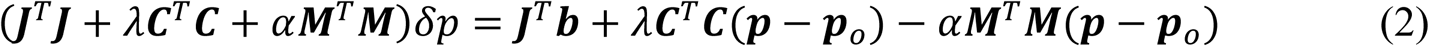

where ***δp*** is the vector of corrections to the initial model parameters, ***p****_j,_* defined as the logarithms of electrical conductivity; ***p****_o_* is the reference model vector, representing prior knowledge or assumptions about the subsurface; ***b*** is the data misfit vector, containing the differences between the logarithms of observed and calculated EC_a_ values; ***J*** is the Jacobian matrix, whose elements represent the sensitivity of *i*-th forward-modelled EC_a_ values to changes in the *j*-th model parameter. The superscript *T* denotes matrix transposition. The Lagrange multiplier *λ* controls the amplitude of parameter corrections in the spatial domain, and ***C*** is the spatial roughness coefficient matrix, defined by the four neighbouring locations (north, south, east and west) at time t. The temporal damping factor α controls the magnitude of parameter corrections in the temporal domain and was set to 0.2 for spatiotemporal inversion. When α = 0, the equation reverts to quasi-2D inversion. ***M*** is a square matrix of temporal dimension with diagonal and sub-diagonal elements being 1 or −1.

In this study, a uniform initial model (σ = 200mS/m) was used for both quasi-2D and spatiotemporal inversion. The model included 12 layers, with the bottom boundaries of the first 11 layers at depths of 0.1, 0.3, 0.5, 0.7, 0.9, 1.1, 1.3, 1.5, 1.7, 1.9 and 2.1 m, and the 12th layer at infinity. The maximum number of iterations was set to 20. The λ values were set to 0.07 and to values from 0.3 to 2.7 in increments of 0.3.

Different combinations of inversion algorithms (quasi-2D or spatiotemporal), regularisation option (S1 or S2) and λ were evaluated to identify the most suitable σ estimates for predicting soil volumetric water content (θ, cm^3^/cm^3^). For each inversion parametrisation, an artificial neural network (ANN) was trained using ground-truth data from 18 soil cores collected along three transects spanning a gradient in soil water content. The ANN was implemented in JMP Pro (Version 19.0.1) with a single hidden layer comprising three hyperbolic tangent (TanH) activation nodes. Model performance was assessed using five-fold cross-validation to reduce the risk of overfitting. Following calibration, ANN-predicted θ was evaluated against NMM measurements. The inversion parametrisation yielding the highest coefficient of determination (R²) between ANN-predicted θ and NMM measurements was selected for the final analysis.

### 2.4 Deriving sorghum water use and proxy of root traits

To capture continuous spatial dynamics in the predicted θ profiles while modelling systematic variation and uncertainty, tensor cubic smoothing splines were fitted within a linear mixed model framework for each plot and measurement date, following Verbyla et al. (2018). The fitted splines represented θ as a function of depth. Integrals of the cubic smoothing splines are used to derive areas under the curve, following White et al. <u>(1999)</u> and Verbyla and Verbyla <u>(2009)</u>. The area between each fitted θ profile and the crop lower limit obtained from APSoil (No. 7) was used to estimate plant available water (PAW, mm) over the soil profile. In each EMI survey window, given the rainfall amount was negligible (Fig. S2), the difference in the total area between the two θ profiles was divided by the number of days between surveys to estimate mean daily plot-level crop water use (ET, mm/day). Corresponding differences were calculated for each 10 cm soil layer to estimate depth-specific daily crop water use (mm/100mm/day).

The depth of water extraction front (WEF; Fig. 3) was defined as the deepest soil depth where the depth-specific crop water use was zero, corresponding to the intersection of the two θ splines. WEF during flowering (2–8 February) and grain filling (6–19 March) was defined as WEF_flw_ and WEF_mat_, respectively. In the case where the splines did not intersect, WEF was estimated by linear extrapolation using the two deepest predicted θ values. The advance of the water extraction front from flowering to maturity (AEF_flw-mat_) was calculated as the difference between WEF_mat_ and WEF_flw_ (Fig. 3).

**Fig. 3.**
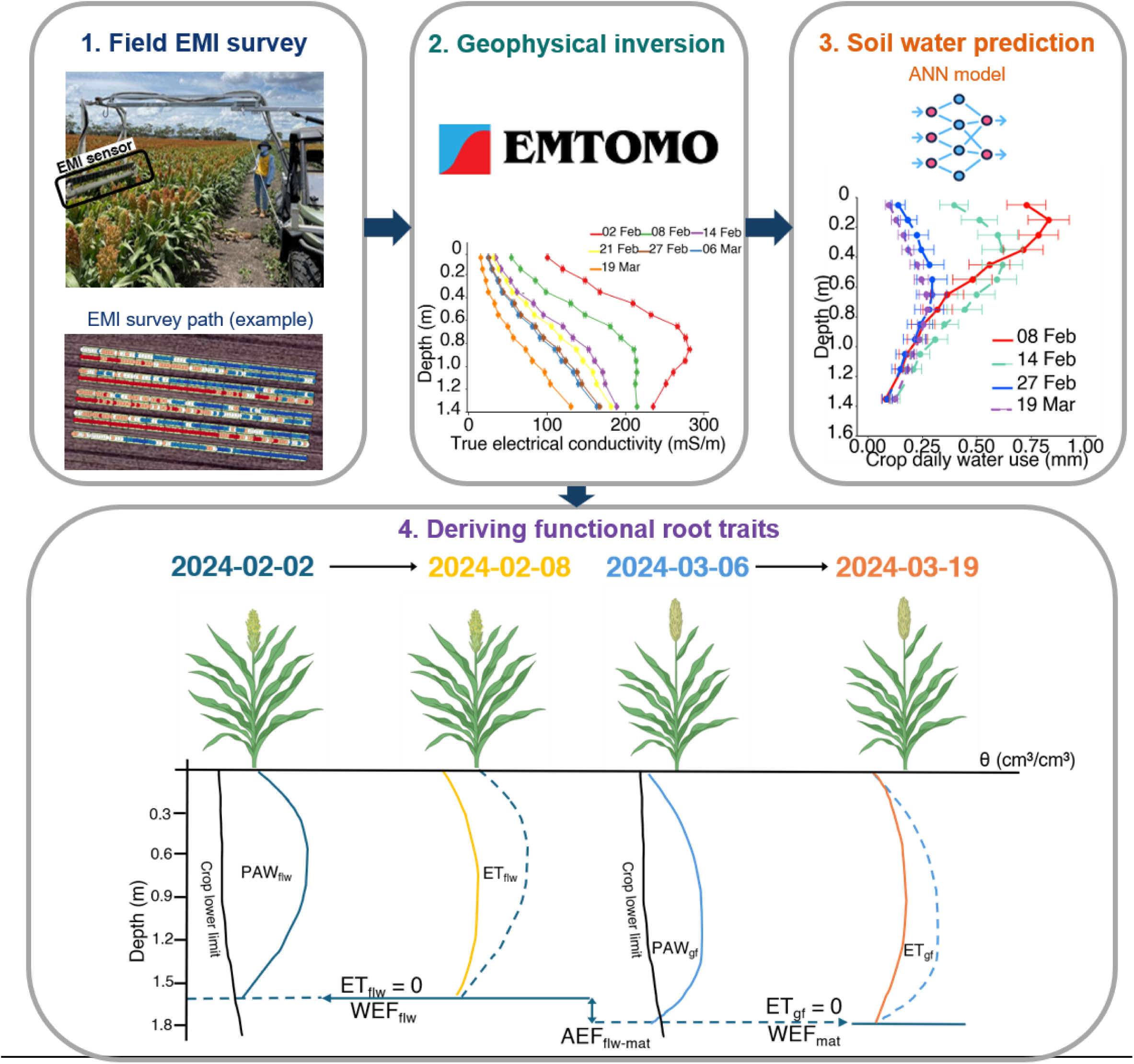
Overview of the EMI-based root function phenotyping workflow. (1) Field EMI survey using the DUALEM-21S sensing system; (2) geophysical inversion to estimate depth-specific true soil electrical conductivity (σ); (3) prediction of soil water profiles using machine-learning algorithms; and (4) schematic derivation of the water extraction front during flowering (WEF_flw_) and grain filling (WEF_mat_) and its advancement from flowering to maturity (AEF_flw-mat_). Blue and yellow curves represent soil volumetric water (θ) profiles from surveys conducted during flowering (2–8 February), and grain filling (6–19 March), respectively. The area between the profiles within each survey window represents crop water use during flowering (ET_flw_) and grain filling (ET_gf_), respectively.

### 2.5 Roots, biomass and yield measurements

To validate the EMI-derived root traits, root coring was conducted for a selected hybrid (K) at maturity. Two soil cores were taken at each sampled plot on the row (5cm from the stem). Cores were divided into 10 cm segments from 0 to 1.2 m depth and 30 cm segments from 1.2 to 2.1 m. Samples were placed in sealed plastic bags and stored in a cold room at 4 °C until root washing using a hydropneumatic elutriation system (Gillison’s Variety Fabrication, Inc). Washed root samples were scanned using an Epson Expression 12000 XL scanner (Seiko Epson Corp., Japan) at 400 dpi. Root length density (RLD, cm/cm^3^) was determined using WinRhizo software.

At flowering, three plants were harvested from a centre row in each plot. Tiller number per plant was counted, and leaf area index (m^2^/m^2^) was measured using an LI-3100C leaf area meter (LI-COR Inc., Lincoln, USA). Shoots were oven-dried at 65 °C to constant weight to determine total aboveground biomass (t/ha). At maturity, eight plants were sampled from uniform areas within the middle two rows of each plot. Shoots were oven-dried at 65 °C to constant weight to determine total maturity aboveground biomass (t/ha). Panicles were threshed to determine grain yield (t/ha), grain number (number/m^2^), and thousand-grain weight (g/10^3^ grains). Grain protein content was determined using near-infrared spectroscopy (Next Instruments Pty Ltd, Australia).

### 2.6 Statistical analysis

Linear mixed models were fitted to response data for yield, yield components and root traits. For above-ground traits, the models included hybrid, row spacing and their interaction as fixed effects. For root traits, the models additionally included depth and all two- and three-way interactions among hybrid, row spacing and depth. Replicates and main-plot terms were included as random effects, and spatial covariance structures were fitted to the residuals (Gilmour et al., 1997). Model parameters were estimated by residual maximum likelihood using the *‘ASReml-R’* package (Butler et al., 2023).

Model assumptions were assessed through visual inspection of residual quantile–quantile plots and plots of residuals against fitted values. No substantial departures from normality or homoscedasticity were observed. Model selection was based on the Akaike information criterion (AIC) and log-likelihood ratio tests, balancing model fit and parsimony. Fixed effects were assessed using Wald tests with approximate F-statistics. Empirical best linear unbiased estimators (eBLUEs) were obtained for significant hybrid, row spacing and interaction effects. The eBLUEs for hybrid by row spacing combinations were used to examine phenotypic relationships among yield, yield components, and root traits. Principal component analysis (PCA) was performed on standardised trait data using the ‘prcomp’ function in *stats* package in R to explore relationships among traits.

To further quantify genetic relationships between the above- and below-ground traits, multi-trait analyses were conducted within a linear mixed model framework. All traits were initially included in the baseline model; however, biomass at flowering and maturity and AEF_flw-mat_ were excluded due to high collinearity caused model convergence failure. The final multi-trait model included flowering leaf area index, grain number, grain weight, ET_flw_, ET_gf_, WEF_flw_ and WEF_mat_ and grain yield. The model included trait, row spacing and their interaction as fixed effects, with hybrid by row spacing interactions fitted as random effects. This allowed estimation of genetic variances for each trait by row spacing combination and genetic covariances between traits within each row spacing. Only traits with significant genetic variance were retained (Table S3 and S4). Genetic covariances were modelled using a factor analytic (FA) variance covariance structure, and the iterative process of fitting FA models is summarised in Table S5. A third-order FA model (FA3) was selected based on the residual maximum-likelihood ratio test and AIC (Table S5). The significance of the genetic covariances was assessed by comparing the FA model with a diagonal model assuming independence between traits through a likelihood ratio test.

The relationship between RLD and water use was modelled using a linear mixed model. Row spacing and RLD were fitted as fixed effects, while replicate, main plot and subplot were included as random design terms. A cubic spline term for RLD was included to accommodate a non-linear relationship between RLD and water use, with a spline by row spacing interaction allowing this relationship to differ among row spacing treatments.

All the above analyses and visualisations were performed in R (R Core Team, 2024).

## 3. Results

### 3.1 Calibration of the EMI data to soil volumetric water content

Fig. 4 shows that the ANN model, calibrated with σ estimates from the inversion modelling, can accurately predict soil volumetric water content (θ, cm^3^/cm^3^). Overall, quasi-2D inversion outperformed spatiotemporal inversion. The largest R^2^ between NMM measurements and ANN-predicted θ was achieved using quasi-2D inversion with the S2 regularisation option, and λ of 0.9 (R^2^ = 0.83; Fig.4a). This configuration was therefore selected for subsequent estimation of crop water-use profiles and derivation of root traits.

**Fig. 4.**
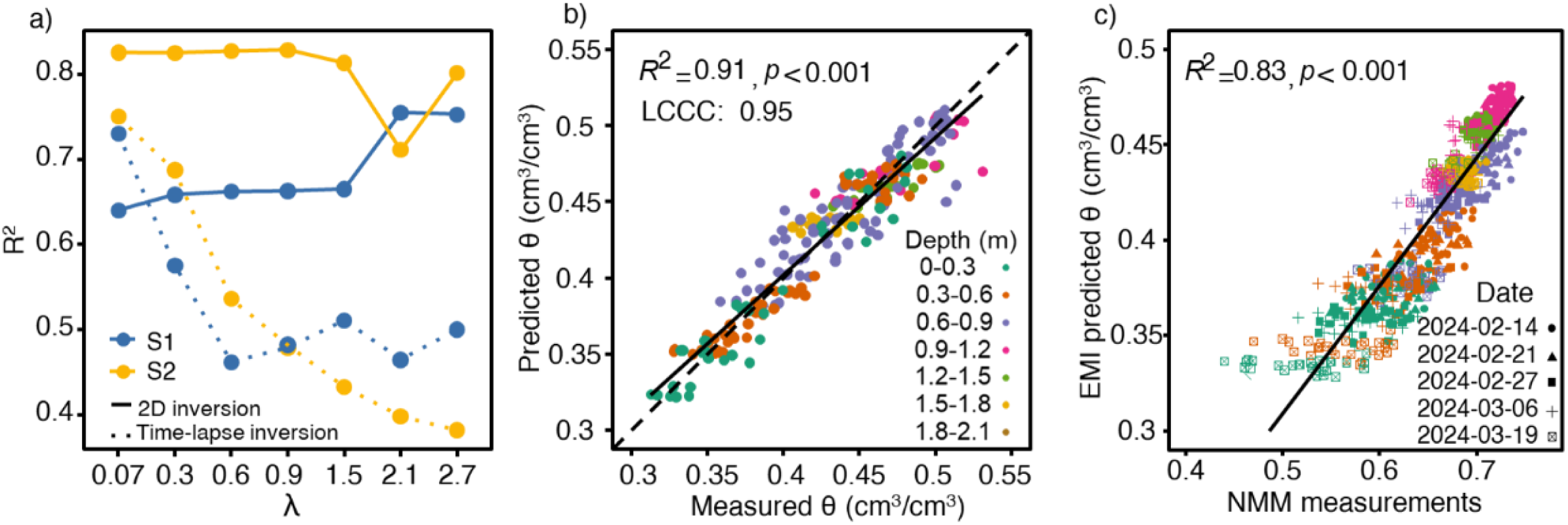
Performance of the artificial neural network (ANN) model for predicting soil volumetric water content (θ, cm^3^/cm^3^) from inverted true soil electrical conductivity. (a) Coefficient of determination (R^2^) between ANN-predicted θ and neutron moisture meter (NMM) measurements as a function of the smoothing factor (λ) for combinations of inversion methods (quasi-2D and spatiotemporal) and regularisation option (S1 or S2); (b) five-fold cross-validation results for the selected model (quasi-2D, S2, λ = 0.9), with the dashed line indicating the 1:1 line; and (c) comparison of ANN-predicted θ and NMM measurements across survey dates.

The corresponding ANN model showed strong predictive performance, with five-fold cross-validation achieving a Lin’s concordance correlation coefficient (LCCC) of 0.95 between predicted and measured θ values (Fig. 4b). The validation against NMM measurements across 27 trial plots and five survey dates yielded an R^2^ of 0.83 (Fig. 4c).

Plant available water (PAW, mm per 100 mm soil) declined drastically across the soil profile over the four EMI surveys as the season progressed (Fig. 5). At flowering, PAW was greatest in the deeper soil layers (0.6-1.2 m). The 0.5 m row spacing generally had lower PAW throughout the soil profile than the 1 m and 1.5 m spacings. This trend was consistent with a deeper WEF_flw_ observed at the 0.5 m spacing, suggesting earlier access to deeper soil water and greater soil water depletion. By maturity (2024-03-19), the PAW profiles were similar among row spacings (Fig. 5).

**Fig. 5.**
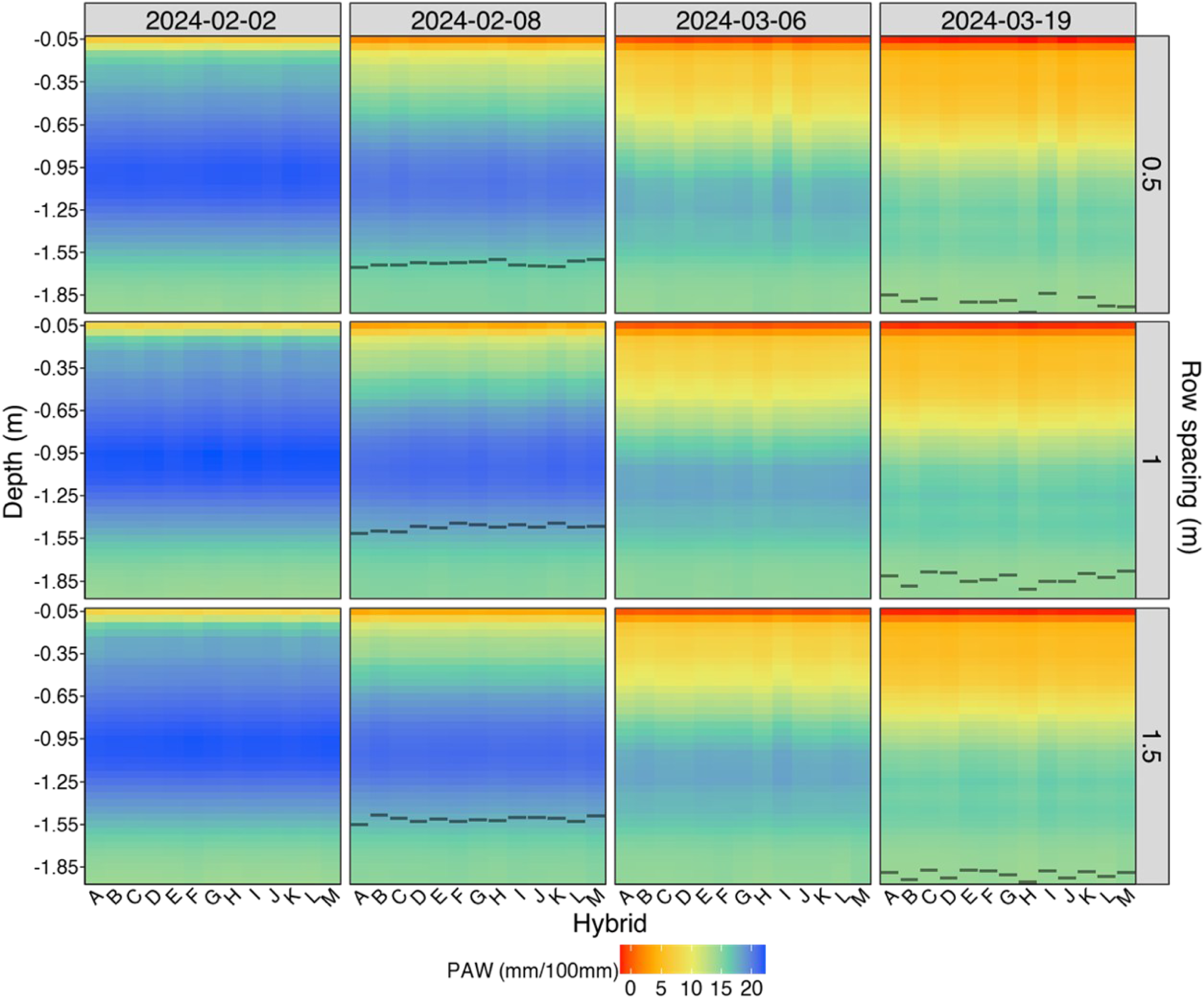
Spatiotemporal distribution of plant available water (PAW; mm per 100 mm soil) for the 13 sorghum hybrids grown at the three row spacings (0.5, 1 and 1.5 m), on the four survey dates (2 and 8 February, and 6 and 19 March). Each panel presents PAW profiles, with hybrids on the x-axis and soil depth (m) on the y-axis. Horizontal lines indicate the water extraction front (WEF) estimated for each hybrid during flowering (2–8 February) and grain filling (6–19 March). Values of PAW and WEF are the empirical best Linear Unbiased Estimators (eBLUEs).

### 3.2 Relationships between the depth-specific crop water use and root length density

Significant relationships were found between EMI-derived layered ET_gf_ and root length density (RLD) for hybrid K at all three row spacings (Fig. 6). The fitted splines indicated a positive correlation between RLD and water use during grain filling, which approached a plateau at shallow soil depths (Fig. 6).

**Fig. 6.**
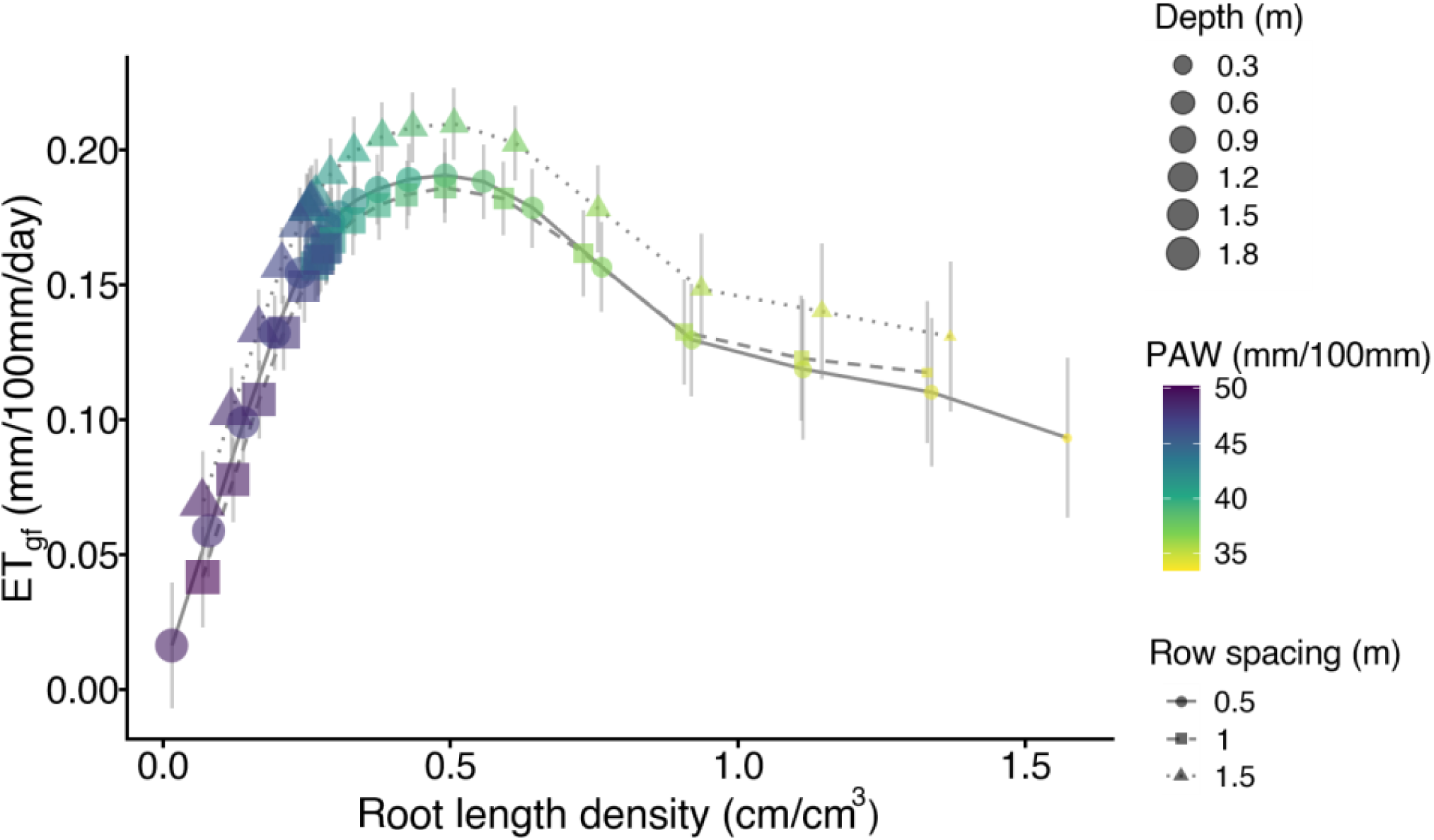
The relationship between root length density (RLD, cm/cm^3^) and depth-specific crop water use during grain filling (ET_gf_) for hybrid K grown at three row spacings (0.5, 1 and 1.5 m). Points represent empirical best Linear Unbiased Estimators (eBLUEs), and error bars indicate standard errors estimated from the linear mixed models. Point shapes and line types represents three row spacings, colours indicate plant-available water (mm/100mm), and sizes represent soil depth (m).

### 3.3 G × M interactions on the root traits

Daily crop water use was higher during flowering (ET_flw_; 4.6-6.6 mm/day) than that during grain filling (2.4-3.6 mm/day) at all three row spacings (Fig. 7; Table 1 and S1). During flowering, the water use was significantly higher at narrower row spacings (*p* < 0.05), with hybrids B and G having the highest water use at the 0.5 m spacing. During grain filling, this pattern reversed, with greater water use observed at wider row spacings. Hybrid B maintained higher water use across all row spacings during this later period (Fig.7 b).

**Fig. 7.**
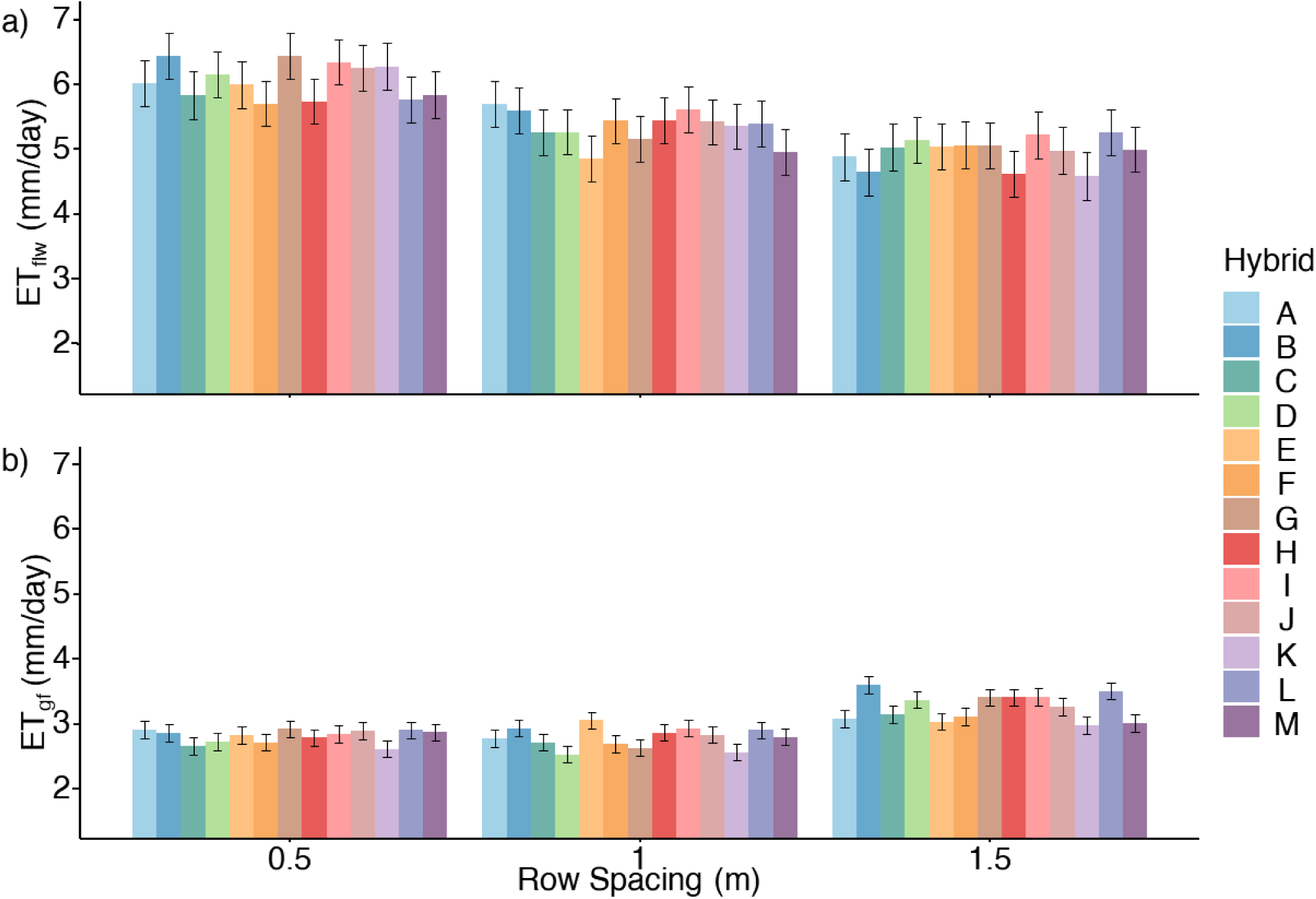
Plot-level daily crop water use (ET, mm/day) during (a) flowering and (b) grain-filling for each hybrid grown under three row spacings. Bars represent empirical best Linear Unbiased Estimators (eBLUEs), and error bars indicate standard errors of the estimations from the linear mixed models.

**Table 1.**
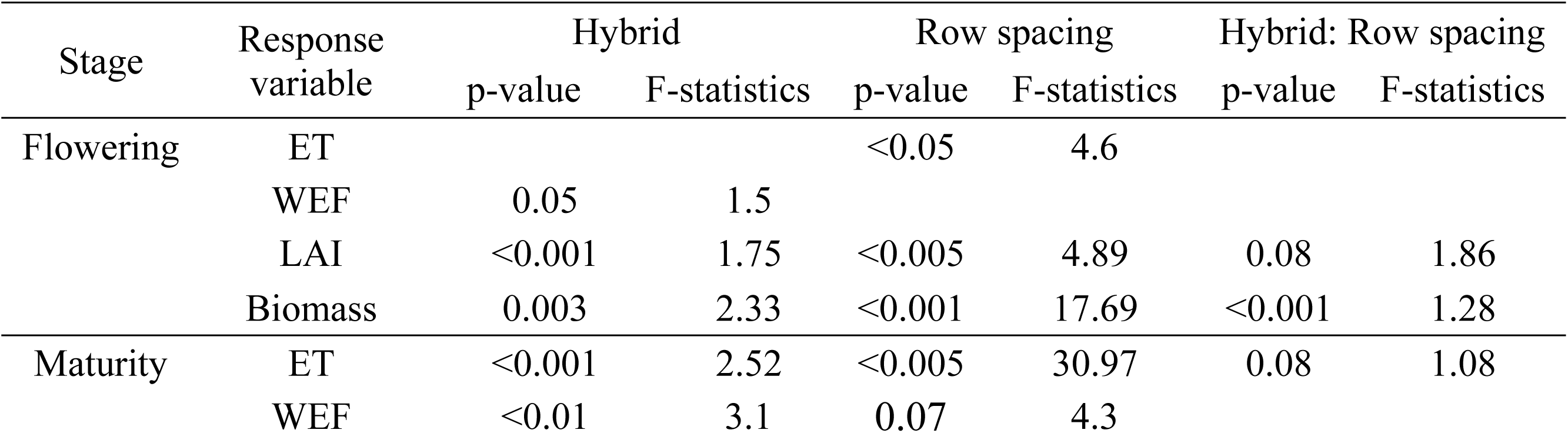
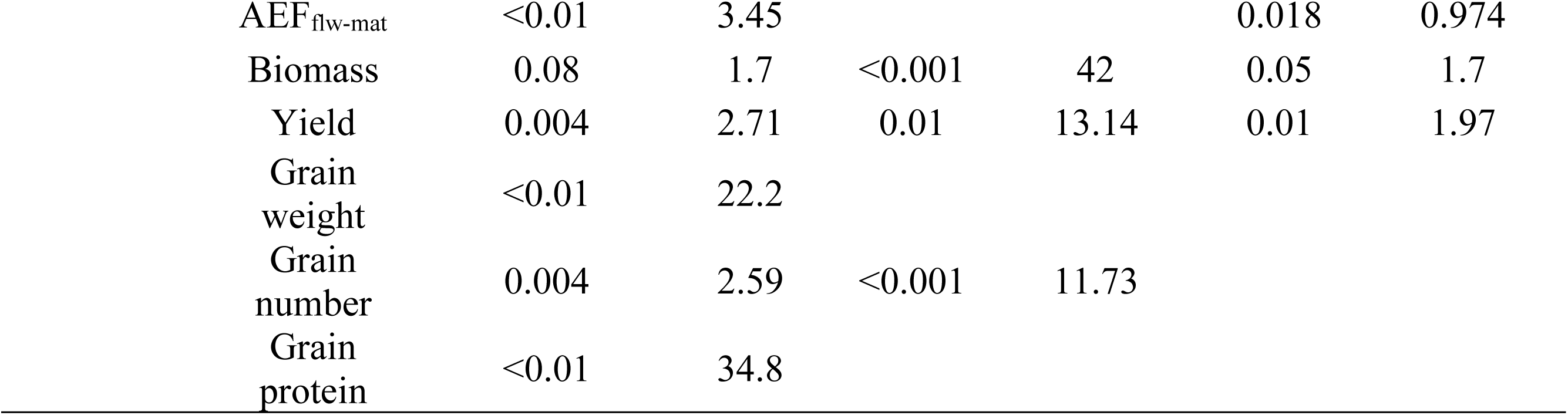
Effects of hybrid, row spacing, and their interaction on evapotranspiration (ET) and water extraction front (WEF) at flowering and during grain filling; WEF advancement from flowering to maturity (AEF_flw-mat_); leaf area index (LAI) and biomass at flowering; and grain weight, grain number, grain yield, biomass, and grain protein content at maturity. Statistical significance was assessed using

| Stage | Response variable | Hybrid |  | Row spacing |  | Hybrid: Row spacing |  |
| --- | --- | --- | --- | --- | --- | --- | --- |
|  |  | p-value | F-statistics | p-value | F-statistics | p-value | F-statistics |
| Flowering | ET |  |  | <0.05 | 4.6 |  |  |
|  | WEF | 0.05 | 1.5 |  |  |  |  |
|  | LAI | <0.001 | 1.75 | <0.005 | 4.89 | 0.08 | 1.86 |
|  | Biomass | 0.003 | 2.33 | <0.001 | 17.69 | <0.001 | 1.28 |
| Maturity | ET | <0.001 | 2.52 | <0.005 | 30.97 | 0.08 | 1.08 |
|  | WEF | <0.01 | 3.1 | 0.07 | 4.3 |  |  |

|  |  |  |  |  |  |  |
| --- | --- | --- | --- | --- | --- | --- |
| AEF<sub>flw-mat</sub> | <0.01 | 3.45 |  |  | 0.018 | 0.974 |
| Biomass | 0.08 | 1.7 | <0.001 | 42 | 0.05 | 1.7 |
| Yield | 0.004 | 2.71 | 0.01 | 13.14 | 0.01 | 1.97 |
| Grain weight | <0.01 | 22.2 |  |  |  |  |
| Grain number | 0.004 | 2.59 | <0.001 | 11.73 |  |  |
| Grain protein | <0.01 | 34.8 |  |  |  |  |

**Table 2.** Empirical best linear unbiased estimates (eBLUEs) for hybrid by row spacing combinations for grain yield (t/ha), biomass at maturity (Biomass_mat_, t/ha), thousand-grain weight (g/10**^3^** grains), grain number (number/m^2^), grain protein content (%), biomass at flowering (Biomass_flw_, t/ha), leaf area index (LAI_flw_, m^2^/m^2^) at flowering, obtained from linear mixed models.

| Hybrid | Biomass<sub>flw</sub> |  |  | LAI<sub>flw</sub> |  |  | Biomass<sub>mat</sub> |  |  | Yield |  |  | Grain weight |  |  | Grain number |  |  | Grain p rotein |  |  |
| --- | --- | --- | --- | --- | --- | --- | --- | --- | --- | --- | --- | --- | --- | --- | --- | --- | --- | --- | --- | --- | --- |
|  | (t/ha) |  |  | (m<sup>2</sup>/m<sup>2</sup>) |  |  | (t/ha) |  |  | (t/ha) |  |  | (g/10<sup>3</sup> grains) |  |  | (number/m<sup>2</sup>) |  |  | (%) |  |  |
| Row spacing | 0.5 | 1 | 1.5 | 0.5 | 1 | 1.5 | 0.5 | 1 | 1.5 | 0.5 | 1 | 1.5 | 0.5 | 1 | 1.5 | 0.5 | 1 | 1.5 | 0.5 | 1 | 1.5 |
| A | 12.2 | 10.3 | 7.8 | 5.2 | 4.6 | 3.1 | 19.9 | 14.4 | 12.6 | 10.2 | 7.3 | 6.7 | 22.5 | 22.7 | 24.1 | 36764 | 36791 | 20960 | 10.5 | 11.2 | 11.1 |
| B | 11.8 | 10.8 | 6.2 | 5.4 | 5.1 | 1.9 | 19.3 | 13.6 | 10.7 | 9.1 | 6.9 | 6 | 24.8 | 25.4 | 26.3 | 27556 | 25476 | 21688 | 10.3 | 10.6 | 9.7 |
| C | 11.9 | 10.3 | 7.5 | 5.7 | 6.4 | 3.0 | 14 | 13.9 | 10.7 | 6.6 | 7.4 | 5.7 | 24.1 | 22.5 | 25.2 | 23636 | 32818 | 25861 | 10.1 | 10.3 | 10.2 |
| D | 9.6 | 9 | 7.1 | 4.5 | 4.0 | 3.8 | 15.1 | 14.6 | 12 | 7.6 | 8.4 | 7 | 21.1 | 21.7 | 24.5 | 32219 | 28297 | 21511 | 9.5 | 9.3 | 9.5 |
| E | 11.6 | 7.7 | 6.6 | 4.3 | 3.8 | 2.9 | 14.5 | 14.7 | 10.8 | 7 | 7.4 | 6.4 | 24.7 | 23.2 | 26.5 | 29938 | 29776 | 20884 | 10.3 | 10.5 | 10.5 |
| F | 12.5 | 9.1 | 10.3 | 6.1 | 4.0 | 4.1 | 14.5 | 12.7 | 11.5 | 6.5 | 6.5 | 5.9 | 26 | 26.4 | 26.7 | 23946 | 26057 | 18804 | 10.7 | 10.4 | 10.8 |
| G | 12.7 | 11.2 | 7.1 | 5.7 | 5.7 | 2.0 | 20.2 | 14.2 | 10.8 | 10.5 | 7.8 | 6 | 23.9 | 22.9 | 22.9 | 32059 | 35225 | 23252 | 10.4 | 10.8 | 10.6 |
| H | 11.8 | 8.8 | 7.7 | 6.4 | 4.0 | 3.6 | 13.9 | 16.3 | 11.3 | 7.3 | 8.1 | 7 | 27.6 | 29.3 | 29.8 | 29099 | 27777 | 28553 | 11 | 11.6 | 11.7 |
| I | 8.7 | 6.5 | 6.3 | 4.1 | 2.8 | 2.5 | 17.2 | 13.4 | 12.1 | 9 | 7.7 | 6.9 | 21.5 | 23.1 | 22.4 | 43699 | 32413 | 27991 | 9.1 | 9.2 | 9.4 |
| J | 8.5 | 9.5 | 9.1 | 3.6 | 4.6 | 4.7 | 15.3 | 12.2 | 11.1 | 8.4 | 6.9 | 5.9 | 27.3 | 27.8 | 26.4 | 21026 | 22685 | 20074 | 11.2 | 11.2 | 11 |
| K | 12.9 | 7.6 | 9.9 | 6.1 | 4.3 | 4.0 | 17.4 | 13.1 | 10.7 | 8.4 | 7 | 5.1 | 28 | 27.9 | 28.3 | 30614 | 29081 | 22436 | 10.6 | 10.6 | 10.8 |
| L | 10 | 8.7 | 6 | 5.4 | 3.9 | 2.9 | 17.6 | 15.5 | 12 | 8.8 | 8.6 | 6.8 | 24.5 | 24.3 | 25.5 | 30392 | 22979 | 22178 | 9.6 | 9.9 | 9.5 |
| M | 10.7 | 9.8 | 8.6 | 4.8 | 3.5 | 3.8 | 13 | 13 | 12.6 | 6.2 | 7.3 | 7 | 29.8 | 27.9 | 28.6 | 13592 | 24387 | 20745 | 10.8 | 10.6 | 10.8 |

WEF_flw_ differed significantly among hybrids and row spacings (*p* < 0.05; Table 1). The 0.5 m row spacing generally had a deeper WEF_flw_ (1.62 m) than the 1.0 and 1.5 m row spacings (1.47 and 1.51 m, respectively). By maturity, all three row spacings reached a similar WEF (1.91, 1.83, 1.91 m at 0.5, 1, 1.5 m spacings, respectively) (Fig. 8b; Table 1). AEF_flw-mat_ differed significantly among hybrids, ranging from 0.19 to 0.46 m (*p* < 0.01), indicating variation in their capacity to access deeper soil water after flowering. Wider row spacings showed greater AEF_flw-mat_, averaging 0.36 and 0.39 m at the 1.0 and 1.5 m spacings, respectively, compared with 0.28 m at the 0.5 m spacing.

**Fig. 8.**
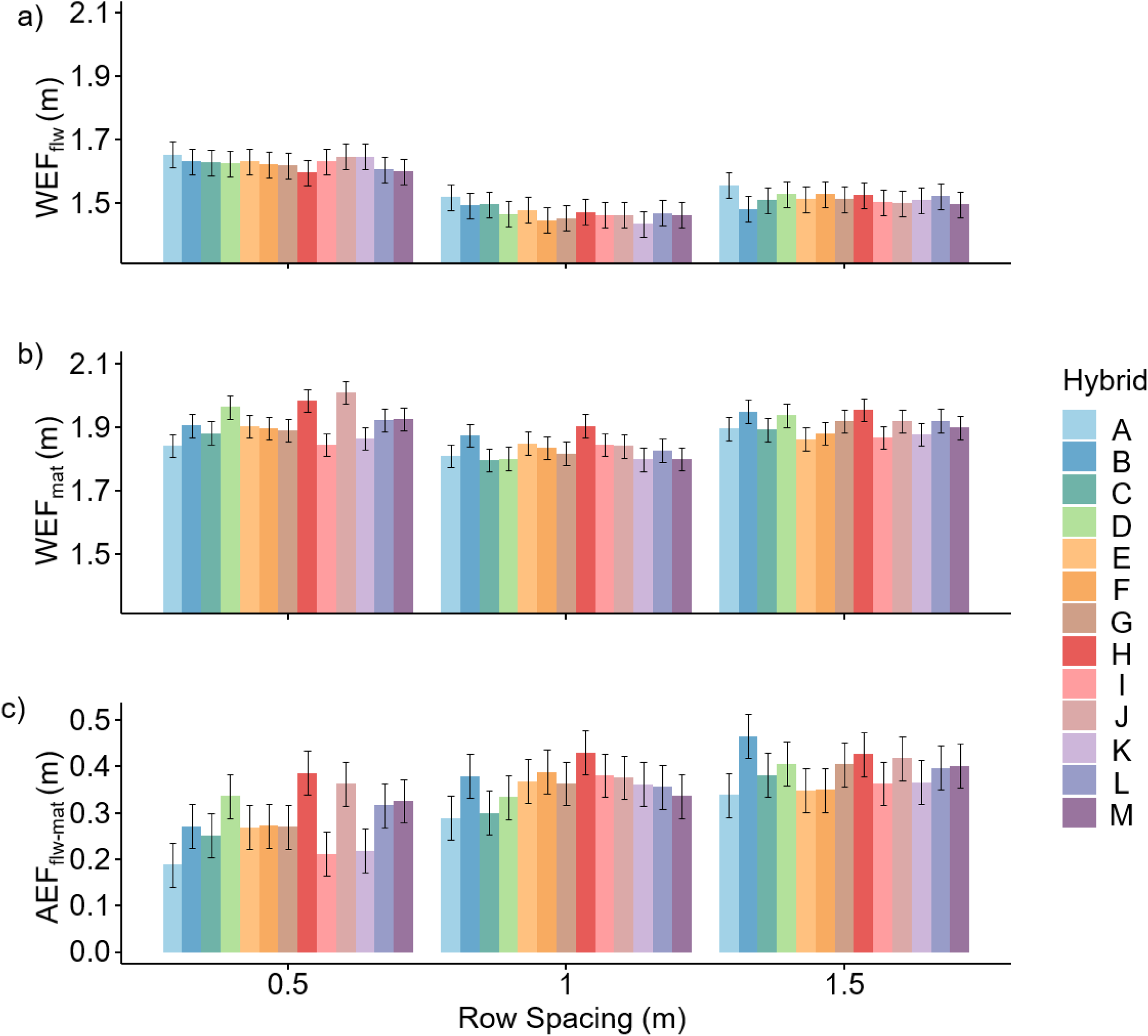
Depth of the water extraction front (WEF, m) during (a) flowering and (b) grain filling, and (c) advancement of the water extraction front (AEF_flw-mat_, m) between flowering and maturity for each hybrid grown at three row spacings. Bars represent empirical best Linear Unbiased Estimators (eBLUEs), and error bars indicate standard errors of the estimations from the linear mixed models.

### 3.4 Canopy traits and yield

The different row spacings exposed the crop to contrasting growing conditions. At flowering, biomass ranged from 6.2 to 12.2 t/ha and LAI from ∼3 to 6.4. Both traits were significantly affected by hybrid and row spacing (Table 1, 2 and S2), with generally greater biomass and LAI in narrower rows. Grain yield ranged from 5.7 to 10.2 t/ha, with a significant hybrid-by-row spacing interaction (*p* = 0.014). Yields were lowest at the wider row spacings, where within-row plant distances were shortest (approximately 11–16 cm). Hybrids D, E and F performed similarly across all row spacings. Hybrids A and G achieved higher yields than the other hybrids in narrower rows, but lower yields in wider rows. Thousand-grain weight, grain number and grain protein content also differed significantly among hybrids (*p* < 0.001).

### 3.5 Associations between traits

#### 3.5.1 Phenotypic correlations

Row spacing was a major driver of variation in canopy, yield, plant available water, and EMI-derived root traits (Fig. 9). The first two principal components (PCs) explained 40% and 24% of the total variance, respectively, with the three row spacings forming distinct groups along PC1. Biomass at flowering (Biomass_flw_) and maturity (Biomass_mat_), LAI_flw_, grain yield, water use at flowering (ET_flw_) and WEF_flw_ clustered together and were associated with the 0.5 m row spacing, indicating positive correlations among these traits. In contrast, water use during grain filling (ET_gf_) and AEF_flw-mat_ formed a separate group associated with the 1.5 m row spacing. Grain weight, grain protein content and WEF_mat_ clustered together and were generally negatively associated with grain number and yield.

**Fig. 9.**
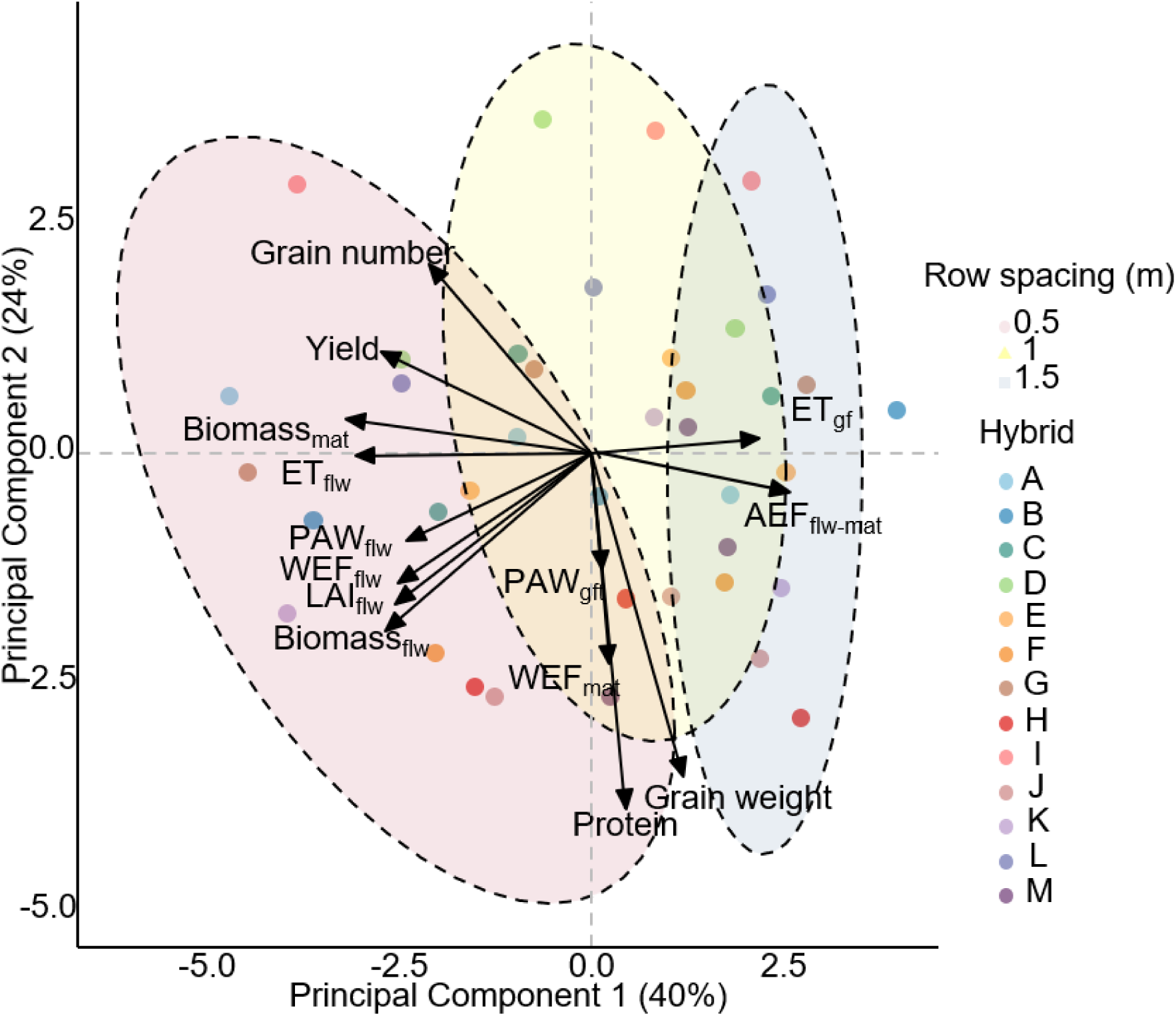
Principal component analysis biplot showing multivariate relationships among yield, canopy, plant available water (PAW) and EMI-derived root traits across row spacings and hybrids. Points represent the 13 hybrids; vectors represent trait loadings, and concentration ellipses indicate groupings by row spacing. Variables include grain yield (t/ha), biomass at maturity (Biomass_mat_, t/ha), grain number (number/m^2^), thousand grain weight (g/10^3^ grains), water extraction front depth during flowering and grain filling (WEF_flw_, WEF_mat_, m), biomass at flowering (Biomass_flw_, t/ha), leaf area index at flowering (LAI_flw_, m^2^/m^2^), grain protein content (%), PAW at flowering and grain filling (PAW_flw_ and PAW_gf_, mm) and daily crop water use during flowering and during grain filling (ET_flw_ and ET_gf_, mm/day).

#### 3.5.2 Genetic correlations

Genetic correlations between traits were estimated using multivariate mixed models that partitioned genetic and non-genetic sources of variance and covariance. Before incorporating all variables into the multivariate model, separate models were fitted for the above- and below-ground traits to retain only traits with significant genetic variance (Tables S3 and S4).

Across all three row spacings, WEF_mat_ was positively correlated with grain weight (r = 0.22 – 0.52; Fig. 10), and there was a consistent trade-off between grain weight and grain number (r = −0.81 to −0.73). The genetic correlation between WEF_mat_ and the rate of crop water use during grain filling (ET_gf_) was significant at wider row spacings (Fig. 10b and c) but not at the narrower row spacing (Fig. 10a).

**Fig. 10.**
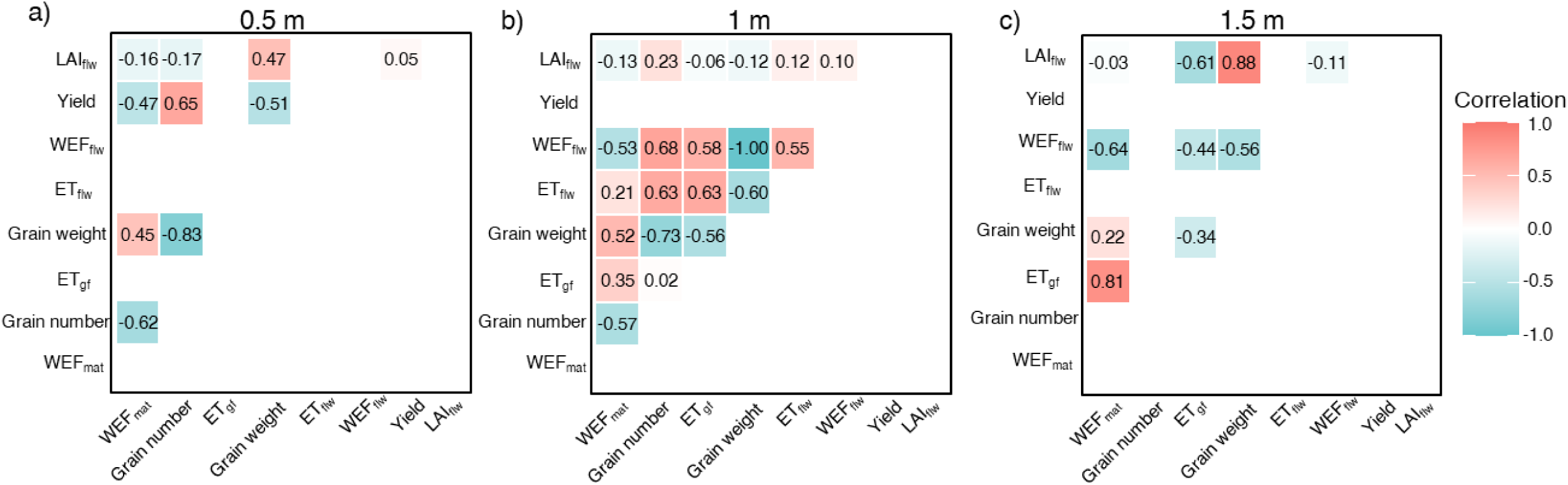
Genetic correlations estimated using a multivariate linear mixed model with a third-order factor analytics (FA3) covariance structure for trait by row spacing combinations with significant genetic variance. Panels show genetic correlations at row spacings of (a) 0.5 m, (b) 1.0 m, and (c) 1.5 m. Colours indicate the direction and magnitude of genetic correlations.

## 4. Discussion

While most root phenotyping approaches focus on root system form, our approach characterises root function quantified through spatiotemporal patterns of crop water use. Within this root functional phenotyping approach, two inversion algorithms were compared (a quasi-2D inversion and a spatiotemporal inversion) to improve soil water prediction accuracy from EMI measurements in a sorghum genotype × management field trial. The non-destructive, high-throughput approach captured spatial and temporal variation in soil-water dynamics at field scale. The quasi-2D inversion provided higher accuracy than the spatiotemporal inversion following optimisation and independent validation. The resulting soil-water profiles enabled derivation of functional root traits associated with yield and yield components.

### 4.1. Non-invasive phenotyping of root functions in the field

Non-destructive, repeatable, high-throughput root phenotyping approaches are essential for both breeding and agronomy. Several non-destructive methods have been proposed; for example, LEADER (Leaf Element Accumulation from DEep Roots) estimates rooting depth from leaf elemental accumulation (e.g., Sr, Si, P, K, Ca, Ba, etc.) (Hanlon et al., 2023). However, the correlation and predictability are site- and species-specific. Ground-penetrating radar (GPR) has been used to detect tree roots with diameters larger than 2mm by emitting electromagnetic pulses (Butnor et al., 2012). Though the method is irrelevant for most grain crops, given that most of the root system is of less than 2mm diameter. Another non-invasive root phenotyping approach is time-lapse soil water monitoring. This approach does not require the individual roots to be spatially resolved, unlike GPR, as it assesses root function by detecting the changes in soil-water dynamics that occur as a result of crop water uptake. Based on this principle, time-lapse EMI surveys, combined with crop ecophysiology principles, have been proposed for the rapid characterisation of root water uptake across both spatial and temporal dimensions in the field (Zhao et al., 2022, 2024, 2025). In their work, indices of soil-layered root activity were closely related to variation in root length density with soil depth across rain-fed and irrigation treatments. Similar results were found here (Fig. 6), supporting the fact that water uptake and root length density are directly associated throughout the profile where root length density is smaller than 1 cm/cm^3^. The plateauing of the relationship indicated that root water uptake was limited by water supply in the drier shallow soil layers or the presence of larger lignified anchoring roots. The consistent relationships between RLD and depth-resolved water uptake indicate that EMI-derived water use was associated with root presence in the soil profile. Notably, the slope of this relationship differs across row spacings. This was also reported previously in Zhao et al. <u>(2024)</u>, with steeper slopes observed under rainfed conditions than under supplementary irrigation. Taken together, these results highlight the importance of phenotyping approaches that capture water uptake dynamics rather than root architecture alone. Although demonstrated here in sorghum, this framework is transferable to other field crops because it is based on crop-induced changes in soil water rather than species-specific root characteristics. Its application to other crops, such as maize, canola, and pulse crops, has significant potential, while site-specific calibration will be needed.

### 4.2. Fine-tuning EMI inversion parameters for estimation of changes in soil water

Inversion of EC_a_ data is used to estimate depth-specific true soil electrical conductivity (σ) profiles. EM4Soil provides multiple inversion algorithms and parameter options for this purpose <u>(Monteiro Santos, 2004)</u>. Using EM4Soil, we systematically compared quasi-2D and spatiotemporal inversion algorithms under different regularisation options (S1 and S2) and smoothing factors (λ). Inversion is inherently non-unique because different σ profiles can correspond to similar EC_a_ responses. We constrained the possible solutions using prior soil-profile information and regularisation. Given that there is no universal best parameter combination, the resulting σ profiles were used to predict soil volumetric water content (θ), and these predictions were subsequently evaluated against independent NMM measurements to determine the most robust inversion configuration.

Our results indicate that quasi-2D inversion consistently outperformed spatiotemporal inversion in both accuracy and stability. The resulting θ predictions showed a consistent positive correlation with NMM measurements across locations and survey dates (Fig. 4c), except in the topsoil layers, where NMM count ratios varied considerably while EMI-derived θ remained relatively stable. The reason could be the inherent limitation of NMM, as measurements near the soil surface can result in neutron escape into the air, and the moisture content could be misestimated (Chanasyk and Naeth, 1996). The poorer performance of the spatiotemporal inversion may reflect the inherent assumption of spatially and temporally smooth variation (Fig. S2). As rainfall causes sudden changes in soil water content that exceed the gradual temporal smoothness assumed by the inversion. Separating surveys into pre- and post-rainfall periods or integrating rainfall as prior information may improve model performance. Differences among parameter combinations also suggest that the inversion remained insufficiently constrained, likely because the sensor measurements provided limited information for resolving conductivity across 12 soil depths (Monteiro Santos et al., 2011). Future improvements may include lifting the sensor to different heights to add additional samples to the inversion’s solution.

### 4.3. Informing G×M effects on the relationship between root function, canopy and yield traits

Understanding the complexity of G×M interactions, alongside the physiological mechanisms underpinning crop responses, can inform agronomic practice and breeding targets. Here, row spacing was used as a management factor to create three environments of contrasting intraspecific competition (Godsey et al., 2012). Differences among the three row spacings in this study reflect the integrated effects of distinct capacities for early carbon acquisition, water use and carbon partitioning efficiency (Lambers, 1983). Early in the season, interception of solar radiation was the main growth-limiting factor (Fig. S2). Under these conditions, the greater distance between plants on the row in the narrower row spacings resulted in reduced intraspecific competition, with the crop achieving full canopy cover and linear growth rates earlier than the 1 and 1.5m row plots, see LAI values at flowering in Table 3. Similar effects of row configurations were observed elsewhere <u>(Godsey et al., 2012)</u>. Fig. 9 also showed a positive association between canopy size and WEF_flw,_ indicating that the deeper water extraction front was most likely driven by the larger crop biomass at flowering.

Here we postulate that wider row spacings (1 and 1.5 m), delayed canopy closure resulted in a smaller canopy by flowering, shallower WEF_flw_, a smaller grain number and grain yield (Table 3, Fig. 9 and 10). Interestingly, particularly for the wider row configurations, the water extraction front continued to deepen after flowering (Fig. 8), which was associated with higher values of water use (ET_gf_) during grain filling (Fig. 9 and 10). This observation contrasts with the assumptions in most crop simulation models. For example, assumptions in the Agricultural Production Systems Simulator (APSIM) (Holzworth et al., 2018), include that the partitioning of new growth to the root system stops at the end of flowering. Our results instead show that this might be inaccurate as between flowering and maturity the depth of the water extraction front increased by more than 0.4m in some combinations of hybrid and row spacing (Fig. 8c). Actually, the observation should be expected, as is supported by the fact that sorghum is a perennial summer cereal. As such, models that account for root growth continuing after flowering may be required to improve modelling outcomes for grain weight and grain quality, particularly under terminal stress environments, as in our trial (Fig. S2).

In grain cereals, many factors affect final grain weight, particularly the balance between sink size (i.e., grain number set during the critical period around flowering) and assimilate availability during grain filling (Sinclair and Jamieson, 2006). The additional advancement of the water extraction front between flowering and maturity (AEF_flw-mat_) contributed to an increased post-flowering water availability, water use and grain weight (Fig. 9). Under terminal drought conditions, crops rely heavily on soil moisture from deep soil layers to delay senescence and sustain assimilate supply during grain filling, i.e., the stay-green trait. Stay-green in sorghum is a complex trait that is associated with the transfer of crop water use between pre- and post-flowering stages, driven by changes in the dynamics of early canopy development (George-Jaeggli et al., 2017), transpiration efficiency (Borrell et al., 2001), as well as deeper root systems (Mace et al., 2012), resulting in delayed senescence and sustained assimilation, larger grain weights and eventually, more stable yields. The observation that grain weight and ET_gf_ were orthogonal in Fig. 9, and the negative correlations between ET_gf_ and grain weight in Fig. 10 b and c, i.e., 1 and 1.5m row spacings, might indicate that grain number had a more dominant effect on grain weights than a punctual estimation of crop water use during grain filling, and provides focus for new, more detailed research.

The PCA analysis in Fig. 9 shows a clear separation of the three row spacings along PC1. Variation along PC2 is attributable to hybrid effects. These are particularly relevant to breeding programs, as genetic performance is well known to be environment dependent. By untangling the relationships among traits across different row spacings, we were able to separate environmental and genetic responses. At 1 m spacing, negative genetic correlations between early-season water use and grain weight, along with positive associations with final yield, indicate a shift in resource allocation. Early-season water use supports yield formation through grain number but at the cost of individual grain weight at maturity. WEF_mat_ showed a positive relationship with grain weight, suggesting that later-stage water acquisition is consistently associated with grain weight, independent of management (Fig. 10).

### 4.4. Limitations of current study and future directions

Several precautions and limitations must be considered when interpreting these findings and applying the proposed framework. First, the current artificial neural network (ANN) model was calibrated using 18 ground-truth soil cores at a fixed sensor height; therefore, direct transfer of this calibration to other pedological settings or alternative sensor configurations is not valid. Application of the method to additional sites requires prior knowledge of the vertical conductivity distribution, thorough comparison of inversion parameters, and local calibration supported by ground-truth data and independent validation. Second, the derived traits reflect root function rather than root architecture. In this study, root length density was measured at maturity, providing a basis for comparison and evidence of correlation, but not validation of temporally resolved patterns of root activity inferred from repeated EMI surveys.

Further, attributing observed soil moisture changes entirely to crop water uptake relies on the assumption of negligible non-transpirational flux. Although surveying during rain-free periods minimized confounding effects and deep drainage was negligible in the heavy-clay profile, surface evaporation and lateral flux cannot be entirely excluded. Finally, the experiment was conducted at a single location, during one growing season, and on heavy clay vertosols. The present findings regarding associations between root traits and yield formation should be regarded as proof-of-concept evidence for incorporating root traits into crop improvement programs. Multi-environment trials across contrasting soil types, seasons, and drought patterns are required to establish a comprehensive physiological understanding of the contributions of root traits to yield and yield components.

Another limitation is the small number of hybrids and treatments examined, which limits its ability to detect genetic variance and correlations between aboveground and belowground traits. Therefore, the correlations specific to spacing in Fig. 10 should be treated as hypotheses for subsequent potential research. Future trials should include more hybrids across multiple environments, and explicitly model genotype, environment, and G×E effects in a multi-trait framework.

## 5. Conclusion

This study shows that EMI, coupled with geophysical inversion and machine learning algorithms, can predict crop water use accurately and efficiently, while the optimal parameter combination requires empirical validation. Three-dimensional water-use maps and valuable root traits were successfully derived for two critical phenological stages. The approach can capture complex G×M effects across growth stages and varying growing conditions. The EMI-derived root traits and water-use profiles showed strong correlations with final yield and have the potential to complement above-ground sensing in support of crop improvement programmes.

## CRediT authorship contribution statement

**Cheng Qian:** Conceptualisation, Data curation, Formal analysis, Software, Visualisation, Writing - original draft, Writing – review and editing; **Daniel Rodriguez:** Conceptualisation, Funding acquisition, Supervision, Writing - review and editing; **Fernando A. Monteiro Santos:** Methodology, Software, Writing – review and editing; **Alison Kelly:** Methodology, Investigation, Formal analysis, Writing – review & editing; **Peter de Voil**: Software, Supervision, Writing – review and editing; **David Jordan:** Supervision, Writing – review and editing.; **Dongxue Zhao:** Conceptualisation, Data curation, Formal analysis, Funding acquisition, Investigation, Methodology, Project administration, Resources, Software, Supervision, Writing – review and editing.

## Data availability

Data will be made available on request.

## Declaration of Generative AI and AI-assisted technologies in the writing process

No AI or AI-assisted technologies were used in writing this paper.

## Acknowledgments

This research was funded by the Australian Government through the Australian Research Council (ARC, IE230100179) and supported by the Grains Research and Development Corporation (GRDC; UOQ2312-009RTX). Dr Zhao is the recipient of an ARC Early Career Industry Fellowship funded by the Australian Government. Cheng Qian is the recipient of a GRDC Research Top-Up Scholarship (UOQ2601-008RSX). We acknowledge Gentech Seeds Pty Ltd for in-kind contribution to the field trial, and Airborn Insight Pty Ltd for conducting the drone flights. We acknowledge Michael Mumford for his statistical support in experimental design. We also acknowledge the technical and casual staff, particularly Ian Broad and Lyle Bromfield, for their assistance with neutron meter measurements and field sampling.

## References

Borrell, A., Hammer, G., Van Oosterom, E., 2001. Stay-green: A consequence of the balance between supply and demand for nitrogen during grain filling? Ann. Appl. Biol. 138, 91–95. 10.1111/j.1744-7348.2001.tb00088.x

Butnor, J.R., Barton, C., Day, F.P., Johnsen, K.H., Mucciardi, A.N., Schroeder, R., Stover, D.B., 2012. Using Ground-Penetrating Radar to Detect Tree Roots and Estimate Biomass, in: Mancuso, S. (Ed.), Measuring Roots. Springer Berlin Heidelberg, Berlin, Heidelberg, pp. 213–245. 10.1007/978-3-642-22067-8_12

Chanasyk, D.S., Naeth, M.A., 1996. Field measurement of soil moisture using neutron probes. Can. J. Soil Sci. 76, 317–323. 10.4141/cjss96-038

D. G. Butler, B. R. Cullis, A. R. Gilmour, B. J. Gogel, and R. Thompson, 2023. ASReml-R Reference Manual.

FAO (Ed.), 2023. Urbanization, agrifood systems transformation and healthy diets across the rural-urban continuum, The state of food security and nutrition in the world. FAO, Rome. 10.4060/cc3017en

George-Jaeggli, B., Mortlock, M.Y., Borrell, A.K., 2017. Bigger is not always better: Reducing leaf area helps stay-green sorghum use soil water more slowly. Environ. Exp. Bot. 138, 119–129. 10.1016/j.envexpbot.2017.03.002

Gilmour, A.R., Cullis, B.R., Verbyla, Arūnas P., Verbyla, Arunas P., 1997. Accounting for Natural and Extraneous Variation in the Analysis of Field Experiments. J. Agric. Biol. Environ. Stat. 2, 269. 10.2307/1400446

Godsey, C.B., Linneman, J., Bellmer, D., Huhnke, R., 2012. Developing Row Spacing and Planting Density Recommendations for Rainfed Sweet Sorghum Production in the Southern Plains. Agron. J. 104, 280–286. 10.2134/agronj2011.0289

Hanlon, M.T., Brown, K.M., Lynch, J.P., 2023. LEADER (Leaf Element Accumulation from Deep Roots): a nondestructive phenotyping platform to estimate rooting depth in the field. 10.1101/2023.05.02.539074

Holzworth, D., Huth, N.I., Fainges, J., Brown, H., Zurcher, E., Cichota, R., Verrall, S., Herrmann, N.I., Zheng, B., Snow, V., 2018. APSIM next generation: Overcoming challenges in modernising a farming systems model. Environ. Model. Softw. 103, 43–51. 10.1016/j.envsoft.2018.02.002

Huang, J., Monteiro Santos, F.A., Triantafilis, J., 2016. Mapping soil water dynamics and a moving wetting front by spatiotemporal inversion of electromagnetic induction data. Water Resour. Res. 52, 9131–9145. 10.1002/2016WR019330

James Wait, 1982. Geo-Electromagnetism. Elsevier. 10.1016/B978-0-12-730880-7.X5001-7

Kim, K.-J., Cho, I.-K., 2011. Time-lapse inversion of 2D resistivity monitoring data with a spatially varying cross-model constraint. J. Appl. Geophys. 74, 114–122. 10.1016/j.jappgeo.2011.04.010

Lambers, H., 1983. “The functional equilibrium”, nibbling on the edges of a paradigm. Neth. J. Agric. Sci. 31, 305–311. 10.18174/njas.v31i4.16935

Mace, E.S., Singh, V., Van Oosterom, E.J., Hammer, G.L., Hunt, C.H., Jordan, D.R., 2012. QTL for nodal root angle in sorghum (sorghum bicolor L. Moench) co-locate with QTL for traits associated with drought adaptation. Theor. Appl. Genet. 124, 97–109. 10.1007/s00122-011-1690-9

Monteiro Santos, F.A., 2004. 1-D laterally constrained inversion of EM34 profiling data. J. Appl. Geophys. 56, 123–134. 10.1016/j.jappgeo.2004.04.005

Monteiro Santos, F.A., Triantafilis, J., Bruzgulis, K., 2011. A spatially constrained 1D inversion algorithm for quasi-3D conductivity imaging: Application to DUALEM-421 data collected in a riverine plain. GEOPHYSICS 76, B43–B53. 10.1190/1.3537834

Pielou, E.C., 1962. The use of plant-to-neighbour distances for the detection of competition. J. Ecol. 50, 357. 10.2307/2257448

Richard, C., Hickey, L.T., Fletcher, S., Jennings, R., Chenu, K., Christopher, J.T., 2015. High-throughput phenotyping of seminal root traits in wheat. Plant Methods 11, 13. 10.1186/s13007-015-0055-9

Sasaki, Y., 2001. Full 3-D inversion of electromagnetic data on PC. J. Appl. Geophys. 46, 45–54. 10.1016/S0926-9851(00)00038-0

Sasaki, Y., 1989. Two-dimensional joint inversion of magnetotelluric and dipole-dipole resistivity data. Geophysics 54, 254–262. 10.1190/1.1442649

Sinclair, T.R., Jamieson, P.D., 2006. Grain number, wheat yield, and bottling beer: An analysis. Field Crops Res. 98, 60–67. 10.1016/j.fcr.2005.12.006

Van Oosterom, E.J., Borrell, A.K., Deifel, K.S., Hammer, G.L., 2011. Does increased leaf appearance rate enhance adaptation to postanthesis drought stress in sorghum? Crop Sci. 51, 2728–2740. 10.2135/cropsci2011.01.0031

Verbyla, A.P., De Faveri, J., Wilkie, J.D., Lewis, T., 2018. Tensor Cubic Smoothing Splines in Designed Experiments Requiring Residual Modelling. J. Agric. Biol. Environ. Stat. 23, 478–508. 10.1007/s13253-018-0334-9

Verbyla, K.L., Verbyla, A.P., 2009. Estimated breeding values and association mapping for persistency and total milk yield using natural cubic smoothing splines. Genet. Sel. Evol. 41, 48. 10.1186/1297-9686-41-48

Wasson, A.P., Rebetzke, G.J., Kirkegaard, J.A., Christopher, J., Richards, R.A., Watt, M., 2014. Soil coring at multiple field environments can directly quantify variation in deep root traits to select wheat genotypes for breeding. J. Exp. Bot. 65, 6231–6249. 10.1093/jxb/eru250

White, I.M.S., Thompson, R., Brotherstone, S., 1999. Genetic and environmental smoothing of lactation curves with cubic splines. J. Dairy Sci. 82, 632–638. 10.3168/jds.S0022-0302(99)75277-X

Zhao, D., De Voil, P., Sadras, V.O., Palta, J.A., Rodriguez, D., 2025. The plasticity of root traits and their effects on crop yield and yield stability. Plant Soil. 10.1007/s11104-024-07185-6

Zhao, D., deVoil, P., Rognoni, B.G., Wilkus, E., Eyre, J.X., Broad, I., Rodriguez, D., 2024. Sowing summer grain crops early in late winter or spring: effects on root growth, water use, and yield. Plant Soil. 10.1007/s11104-024-06648-0

Zhao, D., Eyre, J.X., Wilkus, E., De Voil, P., Broad, I., Rodriguez, D., 2022. 3D characterization of crop water use and the rooting system in field agronomic research. Comput. Electron. Agric. 202, 107409. 10.1016/j.compag.2022.107409

