## Supplementary file for "Closing the phenotyping gap: non-invasive electromagnetic induction enables high-throughput screening of crop root function in the field"

Supplementary material

A high-throughput phenotyping approach for screening the crop root system in the field

Cheng Qian^1^, et al.

| **Nomenclature** | |
| --- | --- |
| **Abbreviation** | **Definition** |
| EMI | Electromagnetic induction |
| EC_a_ | Apparent soil electrical conductivity |
| σ | Soil true electrical conductivity estimated from inversion |
| θ | Volumetric soil water content |
| WEF_flw_ | Water extraction front at flowering |
| WEF_mat_ | Water extraction front at maturity |
| AEF_flw-mat_ | Advance of water extraction front between flowering and maturity |
| ET_flw_ | Crop evapotranspiration during flowering |
| ET_gf_ | Crop evapotranspiration from grain filling to maturity |
| PAW | Plant available water |
| RLD | Root length density |
| LMM | Linear mixed model |
| FA | Factor analytic |
| eBLUE | Empirical best linear unbiased estimate |
| ANN | Artificial neural network |
| NMM | Neutron moisture meter |
| AIC | Akaike information criterion |
| G×E×M | Genotype × environment × management interaction |

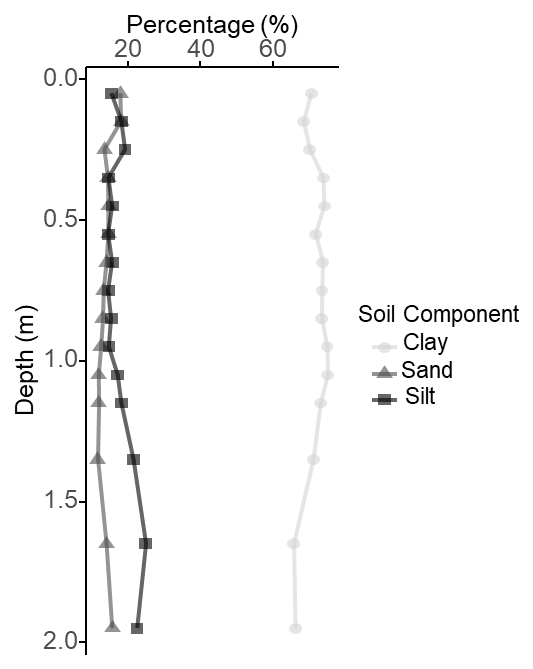

**Fig. S1** Soil characterization results of Tosari site down to two metres. The circle, triangles and squares represent clay, sand and silt content.

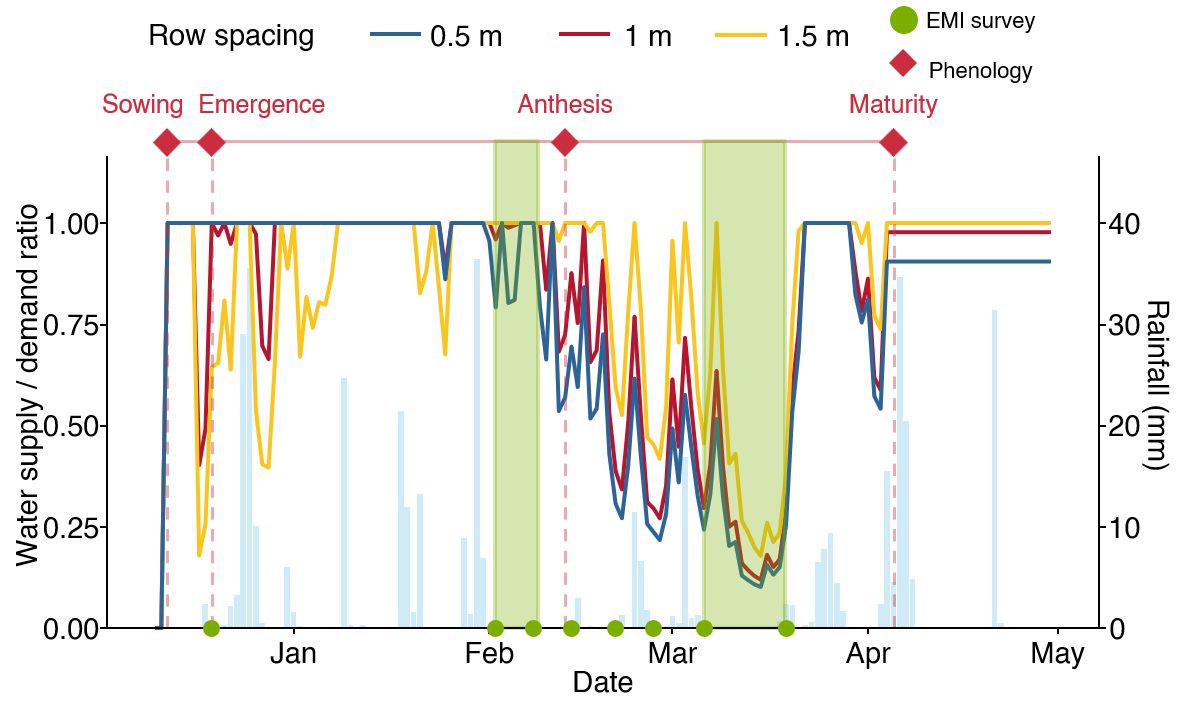

**Fig. S2** APSIM simulated water supply-demand ratio (stress index) for hybrid J with average parameter values. The water supply/demand ratio indicates the degree to which the soil water extractable by the roots (water supply) can match the potential transpiration (water demand). The water demand (mm) corresponds to the amount of water the crop would have transpired without soil water constraint and is estimated daily from the amount of crop growth on that day (g mm^–2^), and the atmospheric saturation vapor pressure deficit (kPa). Line colours correspond to the three row spacings, and the dashed vertical line is the time of flowering.

| **Table S1.** Estimated variance components for random effects at different experimental-design strata for ET and WEF at flowering and maturity, and WEF advancement between flowering and maturity. | | | | | |
| --- | --- | --- | --- | --- | --- |
| Response variable | Model component | Variance component | SE | Z-ratio | Status |
| ET_flw_ | Rep | 5.50E-07 | NA | NA | B |
|  | Mainplot | 0.13 | 0.09 | 1.43 | P |
|  | Plot | 0.34 | 0.05 | 6 | P |
| ET_mat_ | Rep | 0.09 | 0.01 | 0.8 | P |
|  | Mainplot | 0.003 | 0.005 | 0.64 | P |
|  | Plot | 5.00E-02 | 8.00E-03 | 6 | P |
| WEF_flw_ | Rep | 9.00E-04 | 0.002 | 0.46 | P |
|  | Mainplot | 0.003 | 0.002 | 1.37 | P |
|  | Plot | 0.001 | 0.0001 | 6 | P |
| WEF_mat_ | Rep | 4.30E-09 | NA | NA | B |
|  | Mainplot | 1.20E-03 | 8.00E-04 | 1.5 | P |
|  | Plot | 2.70E-03 | 4.00E-04 | 6 | P |
| AEF_flw-mat_ | Rep | 8.00E-04 | 0.002 | 0.43 | P |
|  | Mainplot | 0.002 | 0.002 | 1.27 | P |
|  | Plot | 0.003 | 0.0005 | 6 | P |
| Variance components were estimated using residual maximum likelihood (REML). SE, standard error; P, positive definite; B, bounded. Main-plot and plot-residual terms represent the corresponding experimental-error strata. Z-ratios are presented descriptively as the ratio of the variance-component estimate to its standard error. | | | | | |

| **Table S2.** Estimated variance components for random effects at different experimental-design strata for grain yield, biomass at maturity, grain weight, grain number, grain protein content, biomass at flowering, and leaf area index at flowering. | | | | | |
| --- | --- | --- | --- | --- | --- |
| Response variable | Model component | Variance component | SE | Z-ratio | Status |
| Biomass at Flw | Rep | 0.15 | 0.34 | 0.43 | P |
|  | Mainplot | 0.22 | 0.36 | 0.62 | P |
|  | Plot | 3.69 | 0.61 | 6 | P |
| LAI at Flw | Rep | 8.43 | NA | NA | B |
|  | Mainplot | 8.43 | NA | NA | B |
|  | Plot | 8.33E+07 | 1.30E+07 | 6.24 | P |
| Biomass | Rep | 4.60E-07 | NA | NA | B |
|  | Mainplot | 0.06 | 0.24 | 0.24 | P |
|  | Plot | 4.6 | 0.77 | 6 | P |
| Yield | Rep | 0.04 | 0.1 | 0.38 | P |
|  | Mainplot | 0.07 | 0.1 | 0.71 | P |
|  | Plot | 1.22 | 0.19 | 0.48 | P |
| Grain weight | Rep | 2.00E-07 | NA | NA | B |
|  | Mainplot | 0.15 | 0.18 | 0.78 | P |
|  | Plot | 2 | 0.36 | 5.57 | P |
| Grain number | Rep | 0.23 | 1.64 | 0.14 | P |
|  | Mainplot | 4.40E-06 | NA | NA | B |
|  | Plot | 4.3 | 7.8 | 5.6 | P |
| Protein | Rep | 0.04 | 0.05 | 0.93 | P |
|  | Mainplot | 0.001 | 0.007 | 0.18 | P |
|  | Plot | 0.1 | 0.02 | 5.57 | P |
| Variance components were estimated using residual maximum likelihood (REML). SE, standard error; P, positive definite; B, bounded. Main-plot and plot-residual terms represent the corresponding experimental-error strata. Z-ratios are presented descriptively as the ratio of the variance-component estimate to its standard error. | | | | | |

**Table S3.** Estimated genetic variance components for above-ground traits under different row spacings (0.5 m, 1 m, and 1.5 m) from a multi-trait linear mixed model.

| Variable: Row spacing | Variance component | Standard error | z.ratio | Bound | %ch |
| --- | --- | --- | --- | --- | --- |
| Flowering biomass 0.5m | 0.02 | 0.39 | 0.04 | P | 0.1 |
| Flowering biomass 1m | 0 | NA | NA | B | NA |
| Flowering biomass 1.5m | 0 | NA | NA | B | NA |
| Flowering leaf area 0.5m | 0.19 | 0.14 | 1.39 | P | 0 |
| Flowering leaf area 1m | 0.3 | 0.17 | 1.73 | P | 0 |
| Flowering leaf area 1.5m | 0.15 | 0.11 | 1.32 | P | 0 |
| Maturity Biomass 0.5m | 0.01 | 0.29 | 0.03 | P | 6.9 |
| Maturity Biomass 1m | 0 | NA | NA | B | NA |
| Maturity Biomass 1.5m | 0 | NA | NA | B | NA |
| Yield 0.5m | 0.04 | 0.09 | 0.43 | P | 0.1 |
| Yield 1m | 0 | NA | NA | B | NA |
| Yield 1.5m | 0 | NA | NA | B | NA |
| Grain weight 0.5m | 4.08 | 2 | 2.04 | P | 0.1 |
| Grain weight 1m | 4.88 | 2.29 | 2.13 | P | 0 |
| Grain weight 1.5m | 4.05 | 1.96 | 2.07 | P | 0 |
| Protein 0.5m | 0.27 | 0.13 | 2.16 | P | 0 |
| Protein 1m | 0.34 | 0.15 | 2.21 | P | 0 |
| Protein 1.5m | 0.38 | 0.17 | 2.22 | P | 0 |
| Grain number 0.5m | 0.14 | 0.11 | 1.26 | P | 0 |
| Grain number 1m | 0.07 | 0.09 | 0.83 | P | 0 |
| Grain number 1.5m | 0 | NA | NA | B | NA |

**Table S4.** Estimated genetic variance components for below-ground traits under different row spacings (0.5 m, 1 m, and 1.5 m) from the multi-trait linear mixed model.

| Variable:Row spacing | Variance component | Standard error | z.ratio | Bound | %ch |
| --- | --- | --- | --- | --- | --- |
| WEF_flw_ 0.5m | 0.00 | NA | NA | B | NA |
| WEF_flw_ 1m | 0.00 | 0.00 | 0.73 | P | 0 |
| WEF_flw_ 1.5m | 0.00 | 0.00 | 0.02 | P | 0 |
| PAW_flw_ 0.5m | 2.37 | 1.56 | 1.52 | P | 0 |
| PAW_flw_ 1m | 0.00 | NA | NA | B | NA |
| PAW_flw_ 1.5m | 0.00 | NA | NA | B | NA |
| ET_flw_ 0.5m | 0.00 | NA | NA | B | NA |
| ET_flw_ 1m | 0.03 | 0.03 | 1.10 | P | 0 |
| ET_flw_ 1.5m | 0.00 | NA | NA | B | NA |
| WEF_mat_ 0.5m | 0.02 | 0.01 | 1.90 | P | 0 |
| WEF_mat_ 1m | 0.00 | 0.00 | 0.41 | P | 0 |
| WEF_mat_ 1.5m | 0.00 | NA | NA | B | NA |
| PAW_mat_ 0.5m | 0.00 | NA | NA | B | NA |
| PAW_mat_ 1m | 2.74 | 1.85 | 1.48 | P | 0 |
| PAW_mat_ 1.5m | 0.00 | NA | NA | B | NA |
| ET_mat_ 0.5m | 0.00 | NA | NA | B | NA |
| ET_mat_ 1m | 0.01 | 0.01 | 0.79 | P | 0 |
| ET_mat_ 1.5m | 0.05 | 0.03 | 1.70 | P | 0 |
| AEF between flowering and maturity 0.5m | 0.03 | 0.01 | 1.94 | P | 0 |
| AEF between flowering and maturity 1m | 0.00 | 0.00 | 0.50 | P | 0 |
| AEF between flowering and maturity 1.5m | 0.00 | NA | NA | B | NA |

**Table S5**. Summary of factor analytic (FA) models of increasing order (denoted by FA_k_, where k corresponds to the order of the model fitted) fitted to the random trait by row spacing by hybrid effects. The most parsimonious model is indicated by an asterisk, and related information is in bold text.

| Model | No. variance components^†^ | Log likelihood | AIC^‡^ | Total %VAF^§^ |
| --- | --- | --- | --- | --- |
| Diag | 49 | 370.13 | -642.26 |  |
| FA1 | 101 | 458.26 | -714.55 | 73% |
| FA2 | 125 | 472.93 | -695.86 | 86.10% |
| **FA3*** | **149** | **486.15** | **-674.3** | **91.10%** |
| FA4 | 173 | 494.04 | -642.09 | 97.31% |

^†^ The number of variance components estimated, associated with the random genetic, design and spatial effects in the model

‡AIC, Akaike Information Criterion (Akaike 1973)

§Total %VAF, the percentage of variance accounted for by all factors in the FA model fitted to the random G × E effects
